# Time-based quantitative proteomic and phosphoproteomic analysis of A549-ACE2 cells during SARS-CoV-2 infection

**DOI:** 10.1101/2024.06.20.599898

**Authors:** Fátima Milhano dos Santos, Jorge Vindel, Sergio Ciordia, Victoria Castro, Irene Orera, Urtzi Garaigorta, Pablo Gastaminza, Fernando Corrales

**Affiliations:** Functional Proteomics Laboratory, National Center for Biotechnology (CNB-CSIC), Darwin 3, 28049, Madrid, Spain; Department of Molecular and Cell Biology, National Center for Biotechnology (CNB-CSIC), Darwin 3, 28049, Madrid, Spain; Proteomics Research Core Facility, Instituto Aragonés de Ciencias de la Salud (IACS), 50009 Zaragoza, Spain

## Abstract

The outbreak of COVID-19, a disease caused by severe acute respiratory syndrome coronavirus 2, led to an ongoing pandemic with devastating consequences for the global economy and human health. With the global spread of SARS-CoV-2, multidisciplinary initiatives were launched to explore new diagnostic, therapeutic, and vaccination strategies. From this perspective, proteomics could help to understand the mechanisms associated with SARS-CoV-2 infection and to identify new therapeutic targets for antiviral drug repurposing and/or discovery. A TMT-based quantitative proteomics and phosphoproteomics analysis was performed to study the proteome remodeling of human lung alveolar cells transduced to express human ACE2 (A549-ACE2) after infection with SARS-CoV-2. Targeted PRM analysis was performed to assess the detectability in serum and prognostic value of selected proteins. A total of 6802 proteins and 6428 phospho-sites were identified in A549-ACE2 cells after infection with SARS-CoV-2. Regarding the viral proteome, 8 proteins were differentially expressed after 6 h of infection and reached a steady state after 9 h. In addition, we detected several phosphorylation sites of SARS-CoV-2 proteins, including two novel phosphorylation events at S410 and S416 of the viral nucleoprotein.

**Importance:** The differential proteins here identified revealed that A549-ACE2 cells undergo a time-dependent regulation of essential processes, delineating the precise intervention of the cellular machinery by the viral proteins. From this mechanistic background and by applying machine learning modelling, 29 differential proteins were selected and detected in the serum of COVID-19 patients, 14 of which showed promising prognostic capacity. Targeting these proteins and the protein kinases responsible for the reported phosphorylation changes may provide efficient alternative strategies for the clinical management of COVID-19.

## Introduction

Coronaviruses (CoVs) are enveloped positive single-stranded RNA viruses that infect humans as well as other species, which makes these pathogens a health, veterinary and economic problem. Four genera can be distinguished among the *Coronaviridae* family, alpha and betacoronavirus that exclusively infect mammals and gamma and deltacoronavirus with a wider host tropism. CoVs are zoonotic agents infection by which usually correlates with respiratory and enteric diseases that may develop into a severe syndrome with respiratory complications and lung injury for which scarce therapeutic options are yet available (1). Since the discovery of the first human coronavirus in mid-60s (2, 3) large efforts have been dedicated to understanding the mechanistic principles of CoVs biology and pathogenesis. In particular, the SARS-CoV-2 pandemic urged the scientific community to look for new strategies to combat the severe respiratory syndrome frequently associated with COVID-19 disease. To cope with this need, ambitious research programs were launched in 2019 to investigate the interaction of SARS-CoV-2 with the host cell, which have significantly enhanced our understanding of COVID-19 and the management of patients (4). International initiatives, such as the COVID-19 MS Coalition or the “Global Health” Interdisciplinary Platform (PTI-CSIC) were among the collaborative efforts launched to fight COVID-19 pandemic. In all these endeavors proteomics has emerged as a masterpiece providing unprecedented technical resources to study proteomes in their whole complexity, either biological fluids, cells, or tissues that complement other traditional approaches commonly used in biomedical research. Proteins are the main effectors of most cellular functions and constitute the principal intermediate module to transfer the information encoded in the genome into individual phenotypes. Therefore, understanding the dynamic reprogramming of the proteome upon SARS-CoV-2 infection is essential to understanding the cellular response to the infection, defining early diagnostic and prognostic methods, and developing effective therapeutic interventions. Changes in the serum proteome induced by SARS-CoV-2 infection have been widely investigated as they could provide a valuable readout of driver biological processes of COVID-19 progression and offer new opportunities to monitor and hopefully predict the severity of the disease. Studies using different cohorts and platforms have been reported including mass spectrometry and affinity reagents-based methods such as OLINK (5–9). Despite the expected biological heterogeneity and methodological diversity, a remarkable overlapping of differential proteins was reported, leading to the proposal of severity biomarkers that highlighted regulation of acute-phase response and inflammation, blood coagulation, lung and kidney damage, immune response, and complement cascade. The value of these proteins for the management of COVID-19 patients was further supported by the observation that these up- or down-regulated proteins returned to normal values 100 days after patient discharge (5).

The identification of new molecules with antiviral capacity has deserved increasing attention to facilitate the development of strategies to combat SARS-CoV-2 infection. Since 2002 SARS-CoV outbreak, the anticoronavirus activity of many novel and repurposed molecules has been demonstrated *in vitro* (10). Although antiviral therapy had a limited impact on COVID-19 patients globally, it remains critical for SARS-CoV-2 unvaccinated or immunocompromised individuals, as well as to combat new variants that would escape the immune defenses even on immunocompetent persons. The antiviral strategies to combat SARS-CoV-2 can be grouped into four main streams: first, inhibition of viral proteins such as the polymerase with nucleoside analogs such as Remdesivir (11, 12), Molnupiravir (13, 14) or viral proteases, including PF-00835231 (15, 16). Second, inhibition of virus entry, including antibodies that target the RBD domain of the SARS-CoV-2 spike protein (17). Third, stimulation of the immune response with interferon alone and in combination with other antiviral drugs (17). Fourth, inhibition of hijacking of host machinery is needed for viral replication, frequently through repurposed drugs previously approved for other indications, including host protease inhibitors (18), nucleotide and protein synthesis inhibitors, and endosomal trafficking inhibitors, among others (17). Although the main therapeutic outcome during the pandemic was the development of neutralizing antibodies, alternative antiviral treatments represent a milestone in controlling the progression to severe COVID-19, particularly in patients at risk (19).

Understanding the host-SARS-CoV-2 interaction appears as a priority to discover the mechanisms underlying the viral pathogenesis and to deduce new ways to combat COVID-19. SARS-CoV-2 enters the host cell upon the interaction of the Spike protein (S) with the ACE2 receptor at the cell surface through its receptor binding domain (RBD) (20, 21). The S protein is then processed by the serine protease TMPRSS2 to allow fusion with the cell membrane followed by the penetration of the viral genomic RNA (1), which is translated into polyproteins that are subsequently processed into smaller products (non-structural proteins) by virus-encoded proteases. Extensive remodeling of the cell endoplasmic reticulum (ER) leads to the formation of double-membrane vesicles that host the synthesis of the viral RNAs that are translated into the structural and accessory proteins. Structural proteins and viral genomes assemble into new viral particles in the ER and are transported into the cell surface for exocytosis (22).

In this study, we have investigated the dynamic rewiring of the proteome and phosphoproteome induced by SARS-CoV-2 in the host cell to identify driver proteins of the cellular response to the infection. Ultimately, a selected panel of regulated target proteins has been verified for their suitability to stratify COVID-19 patients according to their disease severity.

## Results

### Identification of regulated proteins in A549-ACE2 after exposure to SARS-CoV-2

To elucidate the mechanisms underlying the response of A549-ACE2 cells to SARS-CoV-2 infection, we have performed a differential proteomic and phosphoproteomic analysis of A549-ACE2 at different times after infection with SARS-CoV-2 (Supplementary Figure 1). A549-ACE cells were inoculated with SARS-CoV-2 in a single cycle infection setup (MOI 1). Total RNA samples were collected to determine the viral load at different times post-infection by RT-qPCR. Figure 1A shows that viral RNA steadily accumulates over time, reaching a plateau between 6 to 9 hours post-inoculation. Consistent with these results, infectivity titers in the supernatants reached maximum values at 16 hours (Figure 1B), coinciding with the time points at which maximal RNA accumulation is achieved. Viral load correlates with a progressive accumulation of N protein in the majority of the cells, reaching over 95% positive cells at 16 hpi, as determined by immunofluorescence microscopy (Figure 1C).

**Figure 1.**
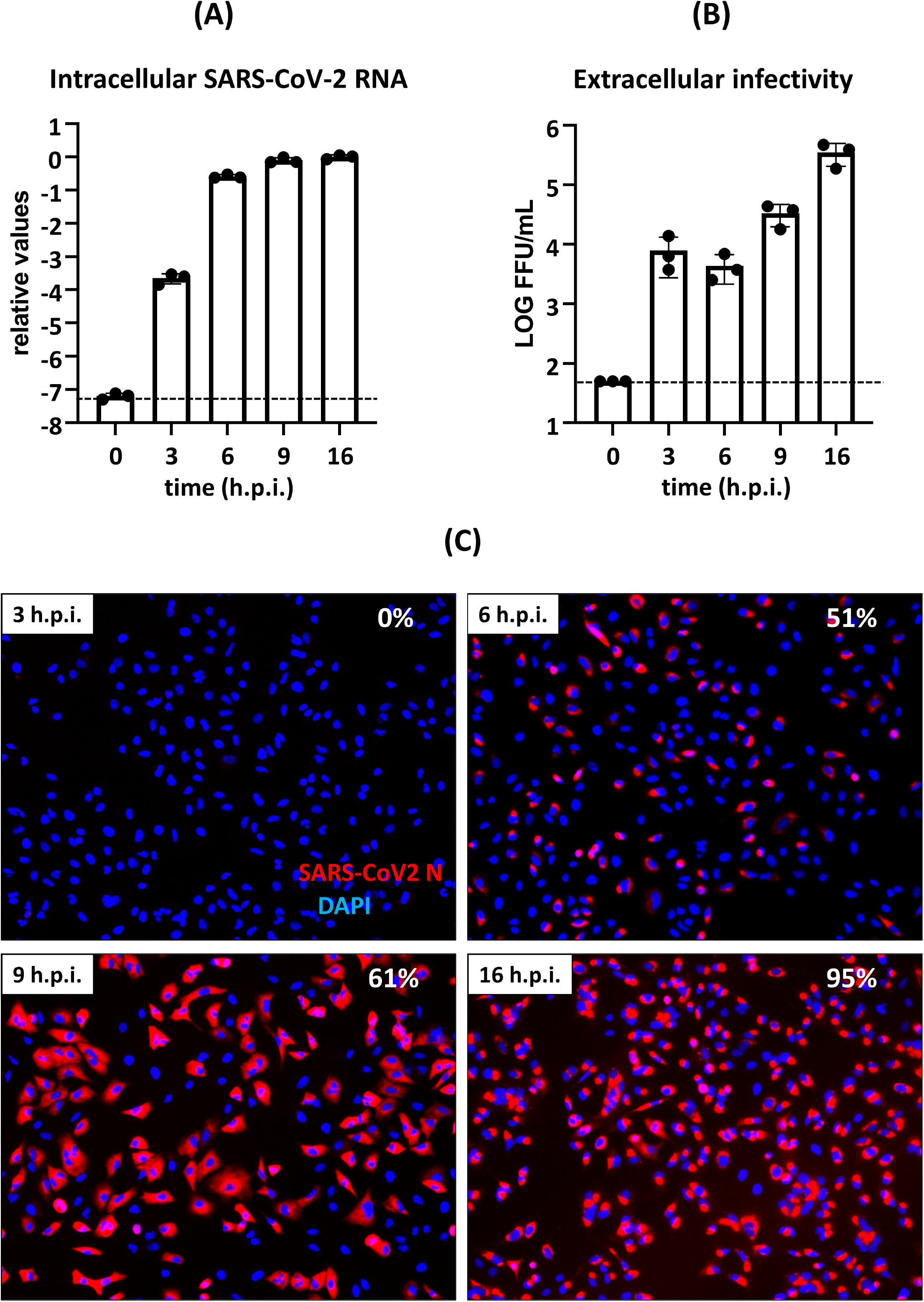
SARS-CoV-2 infection kinetics in A549-ACE2 cells: A549-ACE2 cells were inoculated with SARS-CoV-2 (strain NL/2020; MOI 1). Samples of cells and supernatants were collected at 3, 6, 9 and 16 hours post infection (h.p.i.) to determine relative intracellular genomic RNA levels (panel A) normalised to a cellular housekeeping RNA (28S), extracellular infectivity titers in focus forming units per mL (panel B) and intracellular viral antigen staining (panel C) as described in the Methods section. Data in panels A and B are presented as mean and standard deviation of three biological replicates (n=3). Dotted lines in panels A and B indicate the limit of detection (LOD) determined using mock-infected cell samples as controls (0 hpi). The inset in panel C shows the percentage of N-protein positive cells at each time point.

Altogether, 6,766 protein groups were identified with an FDRΣ1%, from which 6,712 were quantified (Supplementary Table 1), resulting in 91, 115, 261, and 417 regulated proteins (adjusted p-valueΣ5%) after 3, 6, 9, and 16 h of exposure to SARS-CoV-2, respectively (FIG 2A and Supplementary Table 2). Moreover, 10 viral protein groups were also identified, which accumulate over time, except for non-structural protein 8, which reaches a plateau at 9h post-infection (FIG 2C). Upon TiO2 enrichment, 8,223 peptides were identified, corresponding to 2,762 protein groups (FDRΣ1%), revealing 6,428 phosphorylation sites with 100% confidence. Among them, 136, 324, 687, and 902 peptides (adjusted p-value≤5%) displayed a differential pattern at 3, 6, 9, and 16 h post-infection, respectively (FIG 2A and Supplementary Table 2). The overlapping between the panels of regulated proteins by changes in the abundance or phosphorylation is negligible, indicating that the differential phosphorylation events cannot be explained by protein abundance changes (Supplementary Figure 2). Both differential proteins and phosphoproteins efficiently segregated the cell groups at different viral cycle stages, likely reflecting the dynamics of the cellular process involved (FIG 2B). Besides cellular proteins, three SARS-CoV-2 proteins were phosphorylated peaking 6-9 h after cell exposure to the virus, coinciding with maximum replication periods (Figure 1), namely SARS-CoV-2 replicase, membrane protein, and nucleoprotein (FIG 2D). Interestingly, two new phosphorylation sites (S410 and S416) located at the C-terminal peptide of the SARS-CoV-2 nucleoprotein (QLQQSMSSADSTQA) are described here for the first time (Figure 2E and Supplementary Figures 3 and 4).

**Figure 2.**
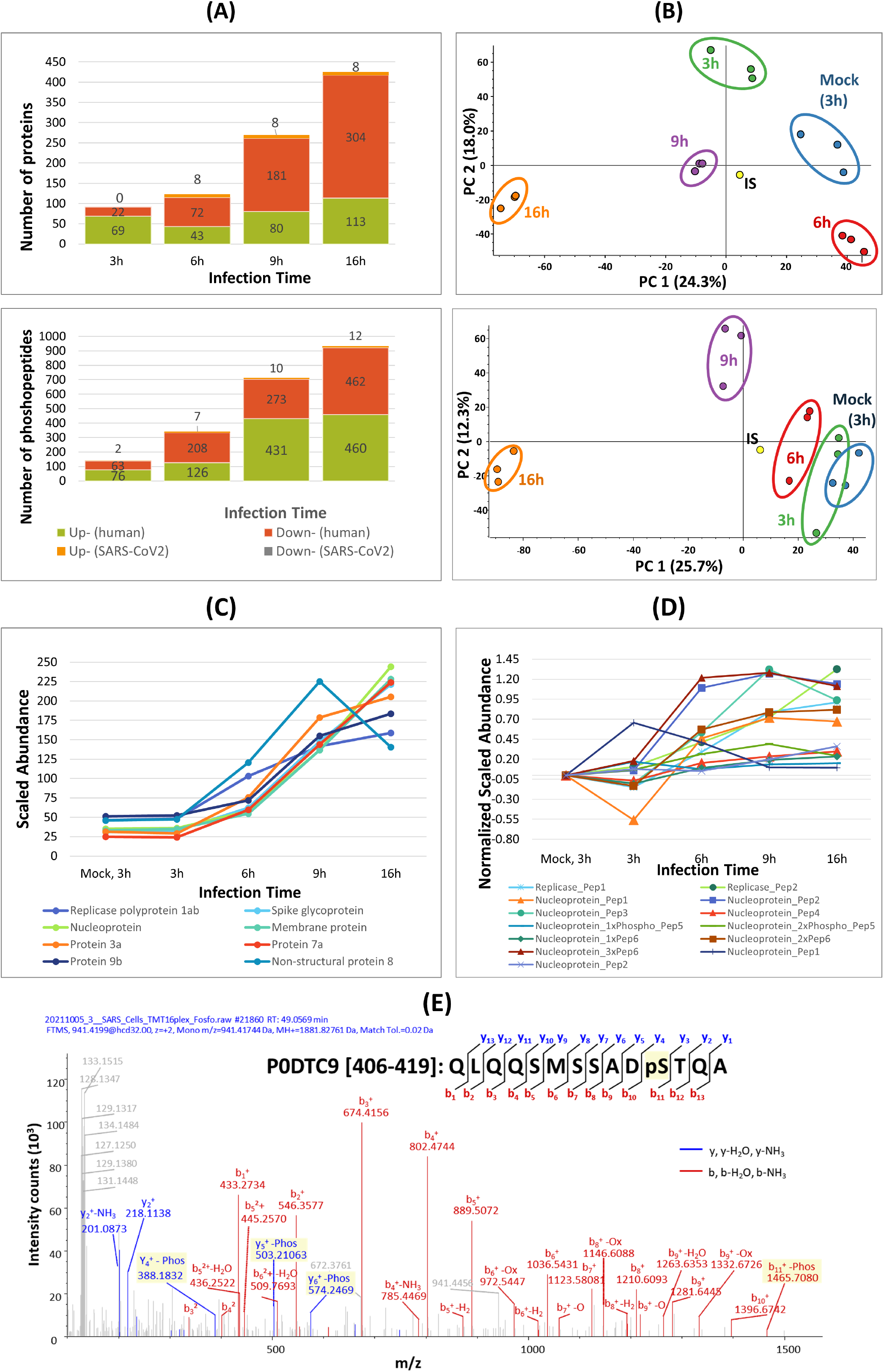
Changes in the host and SARS-CoV-2 proteome and phosphoproteome detected in the analysis of A549-ACE2 cells (n=3) harvested at 3, 6, 9, and 16 h post-infection compared to mock-infected cells (3 h). (A) Number of host (human) and SARS-CoV-2 proteins and phosphopeptides showing a differential pattern at 3, 6, 9 and 16 h post infection. (B) Principal component analysis (PCA) shows that these proteins/phosphopeptides clearly discriminate between different stages of the viral cycle, reflecting host cell dynamics during viral infection. (C) Changes in the levels of SARS-CoV-2 proteins after infection, showing an accumulation of all these proteins over time, except for non-structural protein 8 (D) Phosphorylation dynamics of SARS-CoV-2 proteins including replicase, membrane protein and nucleoprotein during viral infection. (E) Annotated fragmentation spectrum representing a novel phosphorylation site (S416) located on the C-terminal peptide (QLQQSMSSADSTQA) of the SARS-CoV-2 nucleoprotein (P0DTC9 [406-419]).

### Functional analysis of A549-ACE2 differential proteins along SARS-CoV-2 infection

A functional analysis was performed to infer the cellular processes in which the differential proteins and phosphoproteins are involved (Figure 3 and Supplementary Table 3). Overall, formation of the cornified envelope, hemostasis, RNA metabolism and processing, transcription and translation regulation, protein transport and folding, post-translational protein phosphorylation, ribosome biogenesis, cell cycle, signaling (e.g., Rho GTPases), and microtubule/extracellular matrix organization, are the top-rank regulated cellular processes (FIG 3A and 3B). These observations fit well with previous studies that describe how SARS-CoV-2 controls the host cell biology to ensure a successful infection cycle (23–25) (Supplementary Figures 5 and 6). It is also interesting to note that, although not many processes are enriched at the proteomic level, several cellular processes associated with changes in phosphorylation are activated 3h post-infection.

**Figure 3.**
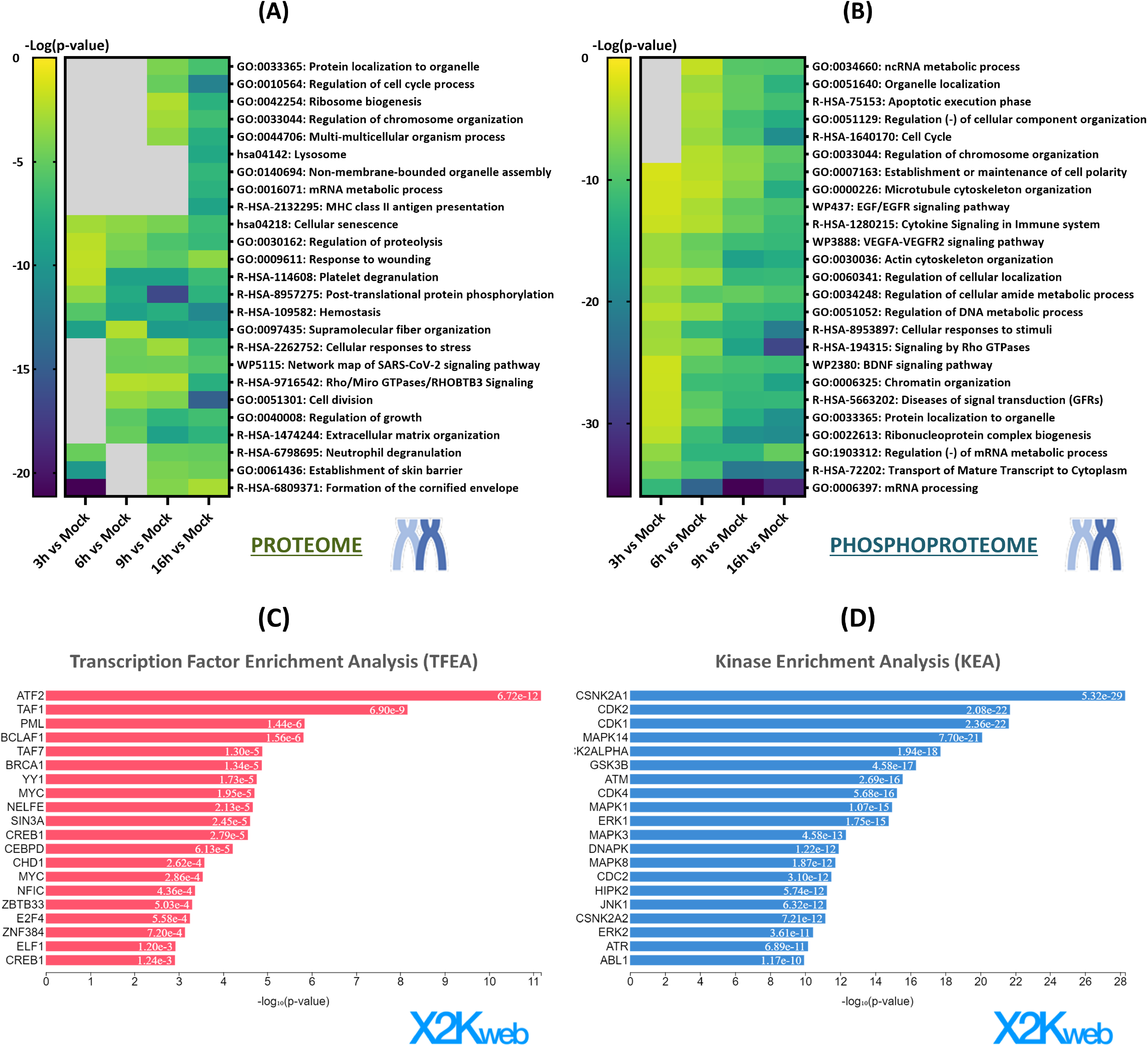
Functional enrichment of the host proteins and phosphoproteins (phosphopeptides) showing a differential pattern upon SARS-CoV-2 infection compared to mock-infected cells (3h). According to the Gene Ontology (GO), Kyoto Encyclopedia of Genes and Genomes (KEGG), Reactome, and WikiPathways databases, heatmaps generated using Metascape display the common and/or unique biological pathways associated with the (A) proteins or (B) phosphopeptides found to be differentially expressed at each infection time point compared to mock-infected cells. (C) Transcription Factor Enrichment Analysis (TFEA) and (D) Kinase Enrichment Analysis (KEA) were performed using the Expression2Kinases webtool of the peptides with differential phosphorylation at 3h post-infection compared to mock-infected cells.

To define upstream regulators that might drive the phosphoproteome changes induced by SARS-CoV-2 in A549-ACE2 cells, a Transcription Factor Enrichment Analysis (TFEA) was used employing the X2Kweb pipeline (Supplementary Figure 7). The analysis suggested an early activation of the transcriptional machinery as expected from the viral requirements for protein synthesis (ATF2, TAF1, YY1, MYC, PML), pointing to the viral control of the cellular apoptotic response through BCLAF1 at 3h post-infection (Figure 3C). Similarly, a Kinase Enrichment Analysis (KEA) of differential phosphopeptides at each time was performed to identify potential kinases whose regulation might lead to the observed changes in protein phosphorylation (Supplementary Figure 7 and Figure 3D). In agreement with previous reports, multiple kinases are activated upon SARS-CoV-2 infection of A549-ACE2 cells (24, 26) which highlights a significant reprogramming of the cellular signaling landscape. Among them, CK2 (CSNK2A1) and CDKs may contribute to the regulation of viral replication (Figure 3D and Supplementary Figure 7).

In order to investigate the dynamics of the cellular processes regulated along the viral infection, the differential proteins and phosphoproteins were first grouped into clusters according to their variation over time. The number of clusters was selected based on their specific time-dependent profile, which was estimated by the minimum distance between cluster centroids (Supplementary Figure 8). According to this criterium, four and six clusters were considered for the differential proteomics (DP clusters) and phosphoproteomics (DPP clusters) analyses, respectively (Figures 4A and 5A). Then, the regulated pathways within each cluster were inferred by functional enrichment analysis (Supplementary Table 4). The top 10 cellular processes regulated by proteins and phosphoproteins of each cluster are shown in Figures 4B and 5B respectively and the specific proteins involved in each case are listed in the Supplementary table 4. DP3 cluster represents proteins that increase transiently at 3h post-infection. Functional analysis revealed an early inflammatory reaction (α2 macroglobulin), deep remodeling of keratin filaments (several keratin species and filaggrin), and response to cellular damage by cell cycle arrest (HSU1). Clusters DP1 and DPP3/6 include proteins showing a steady accumulation or phosphorylation along the infection. As might be expected viral proteins are included in these clusters since they accumulate as the infection cycle progresses until virions are ensembled and released. This has also been verified for the phosphorylation of viral proteins, although the functional outcome of these modifications is not yet clear. Functional enrichment analysis of up-regulated cellular proteins suggests suppression of IFN signaling, as well as cell division and mitosis (NCAPH, CDK1, CDK7, KIF11, BRSK2) and microtubule movement (Kinesines, dynein) Other proteins are involved in cellular process that are known to be regulated by SARS-CoV-2, including autophagy (SQSTM1), histone remodeling (H2A, H2B, H1.2), regulation of Ca^2+^ metabolism (CALM3, CARHSP1), ER stress (KDELR1, ATP13A1, EMC4), and protein folding (PFDN1). Moreover, functional analysis indicates that phosphorylated proteins are involved in the activation of protein import to the nucleus (NUP35, RANBP2, KPNA5, KPNA4, KPNA3, RGPD5), nuclear pore complex assembly, mRNA transport and export from the nucleus (NUP98, NUP35, AHCTF1, RANBP2, RANGAP1, NUP214, NUP153, NUP50, NUP188, ANP32A, NUP107, POM121), activation of translation initiation (EIF4G1, EIF5, EIF5B, CTIF, EIF4B, NCBP1, CSDE1, HSPB1), and microtubule cytoskeleton organization (SON, MAP1B, MAP1S, EML4, DYNC1LI2, MAP4, TACC2, EML1, PHLDB2, CLASP1, TACC3, HAUS6, DYNC1LI1, DST). Clusters DP2 and DP4 integrate down-regulated host cell proteins along the viral cycle. Since the expression dynamic profile is quite similar for proteins in these two clusters, a combined functional analysis was performed considering all downregulated proteins. The most significantly regulated processes include cell adhesion, antigen processing and presentation, response to vitamin D, regulation of IGFR signaling, IL6 signaling, and H2B ubiquitination, which fits well with the phosphorylation of several proteasome proteins (PSMA5, PSMD1, PSMD4, PSMD11, PSMF1). Besides, clusters DPP2 and DPP5 recapitulate proteins whose phosphorylation is reduced as the viral cycle progresses. Similarly to the analysis of down-regulated proteins, the functional enrichment analysis was done by grouping the proteins from both clusters. It revealed that these proteins participate in the regulation of mRNA splicing, immediate filament organization, mRNA processing, adherens junctions organization, rRNA transcription, RNA transport, and regulation of stress fiber assembly. Finally, DDP1 and DDP4 include proteins that transiently decrease or increase respectively their phosphorylation up to 9 h post-infection to then return to basal levels. Regulated pathways by DDP1 cluster proteins included mRNA splicing and processing, H2B ubiquitination, and apoptotic chromosome condensation. Regarding the DDP4 cluster, the principal enriched pathways were the regulation of protein localization and stabilization, microtubule anchoring, actin filament depolarization, and cell-cell adhesion.

**Figure 4.**
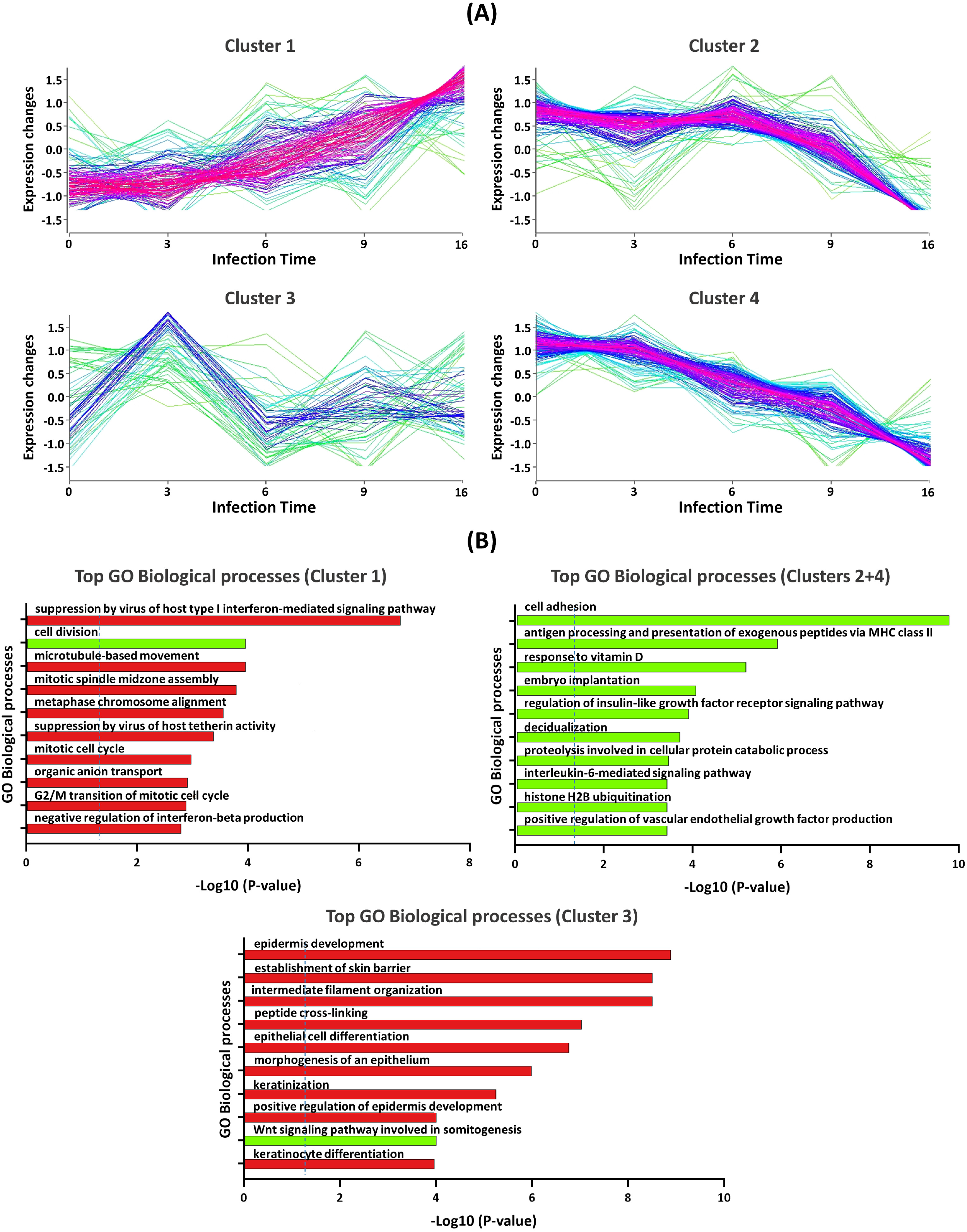
Functional enrichment based on protein expression dynamics in proteome analysis: (A) Mfuzz clustering was applied to define 4 distinct protein clusters with common expression dynamics over the time course of SARS-CoV-2 infection. (B) Functional analysis was performed to assess the top biological processes associated with these Mfuzz-based clusters using the Gene Ontology (GO) database.

**Figure 5.**
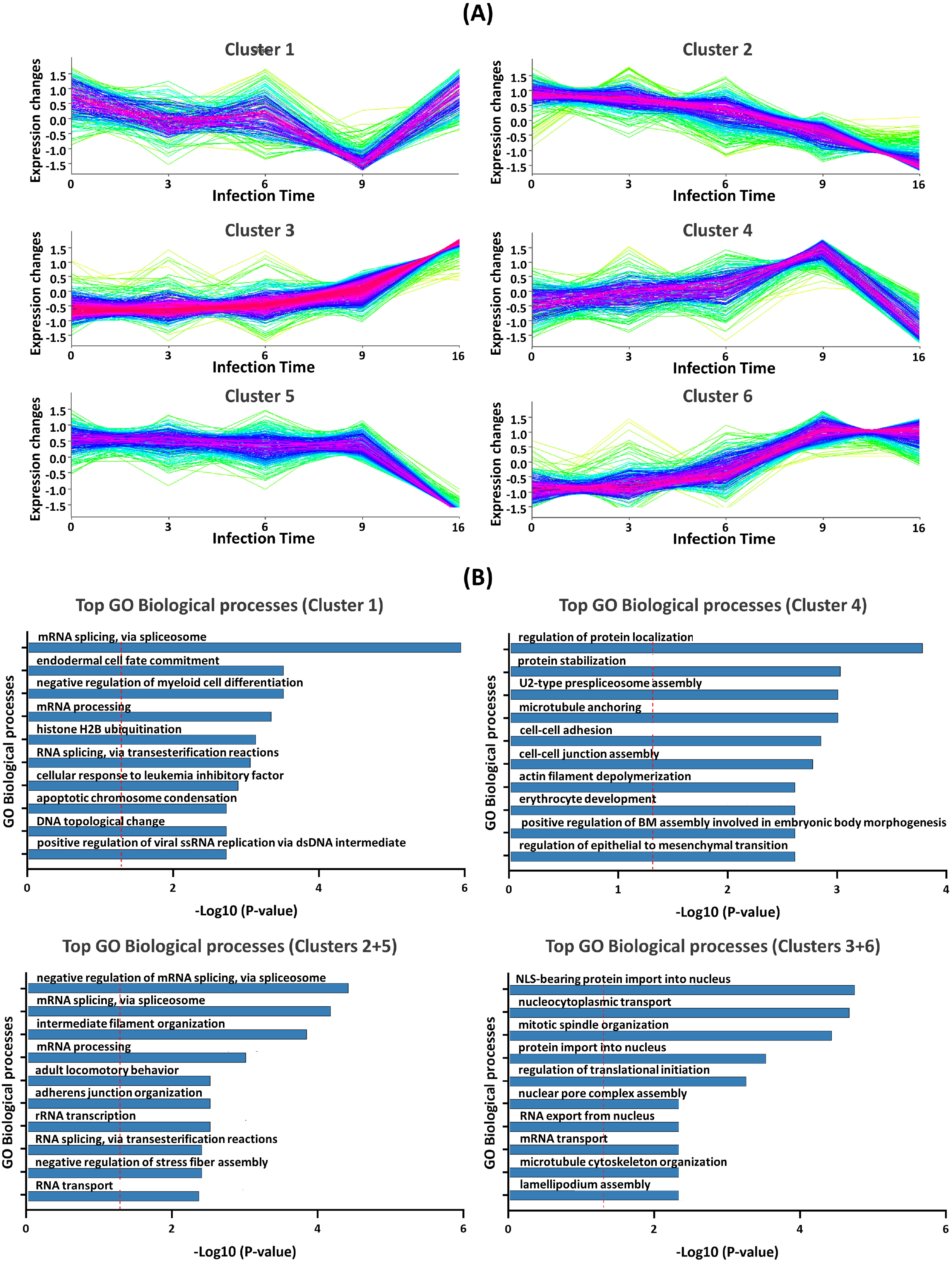
Functional enrichment based on phosphopeptide expression dynamics in phosphoproteome analysis: (A) Mfuzz clustering was applied to define 6 distinct phosphoprotein clusters with common expression dynamics over the time course of SARS-CoV-2 infection. (B) Functional analysis was performed to assess the top biological processes associated with these Mfuzz-based clusters using the Gene Ontology (GO) database.

### Selection of a functionally relevant protein panel and investigation of its value as a plasma readout of COVID-19 severity

Once the mechanisms underlying SARS-CoV-2 infection in lung cells were delineated, it was wondered if the identified driver proteins could be used as a readout of the infection process in humans. To address this question, we focused on proteins among those identified whose presence in serum/plasma has already been demonstrated. The rationale was that these biofluids can be accessed by relatively non-invasive methods, which may facilitate the implementation of the optimized method in future clinical applications. Overall, 826 proteins were selected considering their previous detection in plasma according to Paxdb. Of these, 48 have showed differential expression in cells after exposure to SARS-CoV-2 could be detected in serum/plasma. The top 30 proteins in the importance ranking were then selected by recursive random forest (Supplementary Table 1 and Figure 6). The selected features displayed a time-dependent expression pattern (Figure 6B) that allowed the segregation of the cells along the infection process in the Multidimensional (MDS) plot (3D in Supplementary Figure 9 and 2D in Figure 6A). These proteins represent drivers of the infection that can be detected in plasma/serum and are therefore candidates for the follow-up of COVID-19 patients. Upon optimization and removal of non-detected proteins, 23 proteins were initially quantified by monitoring their proteotypic peptides as indicated in Supplementary Table 5. Desmogelin 1 was not further considered since it could not be detected in most serum samples from COVID-19 patients and, therefore, its quantification was not reliable. The verification of these potential biomarkers was carried out in a cohort of COVID-19 patients with different disease severity, previously characterized in our laboratory (5). This cohort was composed of 76 patients, including 16 non-hospitalized patients (NHOSP), 30 hospitalized patients (HOSP), 10 patients hospitalized in the intensive care unit (ICU), and 20 deceived patients hospitalized in ICU (EXI). Of the 22 proteins successfully quantified (Figure 6C), 14 were found differentially expressed among the different COVID-19 groups (Supplementary Table 6 and Supplementary Figure 10), showing an expression pattern dependent not only on disease severity but also on the age of the patients (Figure 6D).

**Figure 6.**
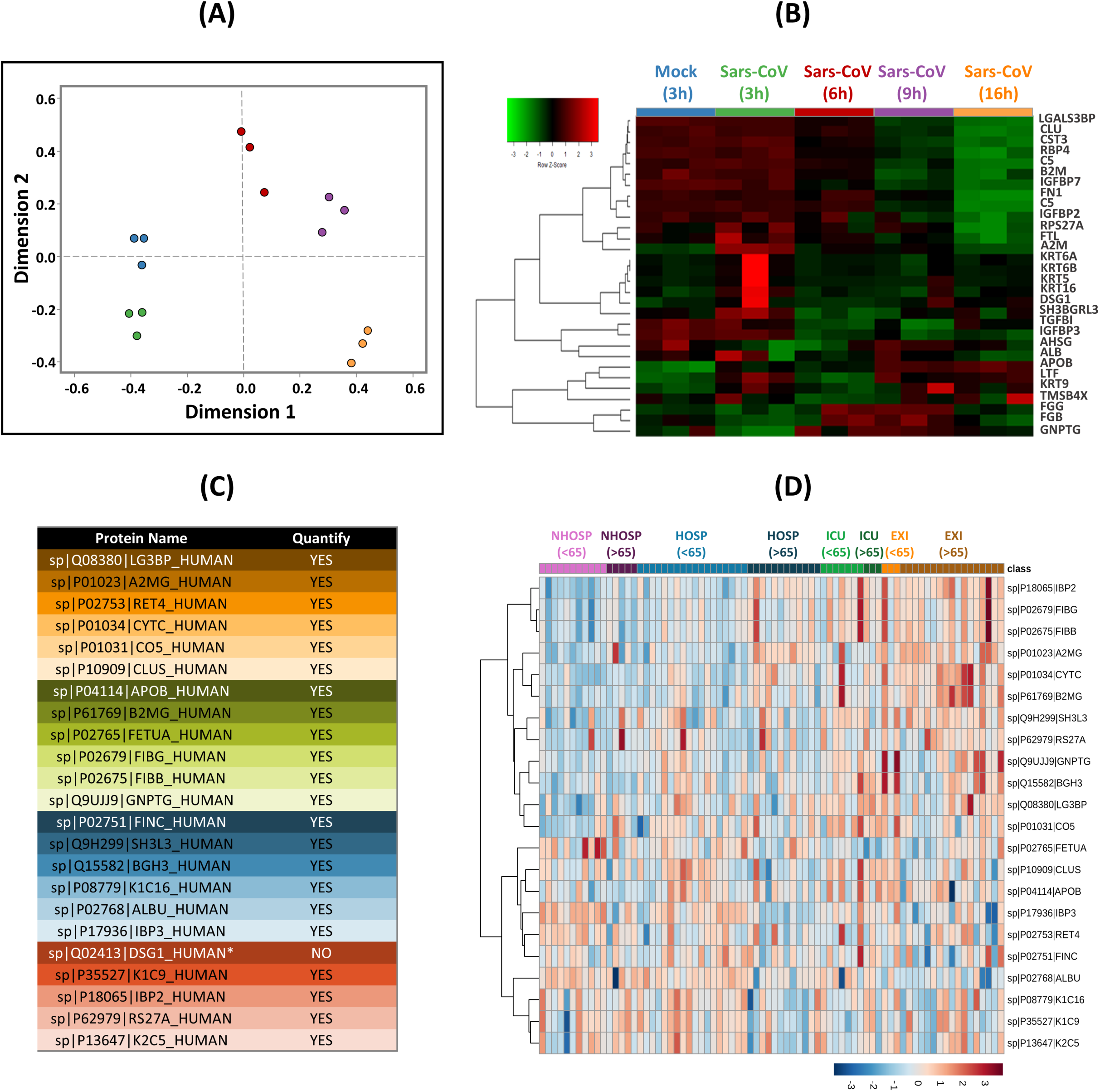
Selection of a functionally relevant protein panel and verification of its value as a plasma readout in the cohort of patients with varying COVID-19 severity. According to the results of the TMT-based proteomics, 30 proteins were selected based on a recursive random forest algorithm and their abundance in human serum/plasma for further validation in serum samples from patients with COVID-19. The selected features (proteins) showed a time-dependent expression pattern in the (B) heatmap, which allowed the segregation of cells along the infection process in the (A) multidimensional (MDS) plot. (C) A list of 22 proteins was successfully quantified by parallel reaction monitoring (PRM). (D) The heatmap shows the abundance of the 14 differentially expressed proteins between the different COVID-19 groups in the PRM analyses, showing an expression pattern that depends not only on the severity of the disease but also on the age of the patients.

### Identification of circulating proteins as readouts of COVID-19 severity using machine learning modeling

Aiming to define a classifier that may help sample stratification, the intensity values extracted from the PRM experiment of the remaining 22 proteins across the 76 COVID-19 serum samples (Supplementary Table 6) were normalized (log2 transformed and scaled by Z-score calculation). The full data set was divided into a training data set consisting of 70 % of the samples, and the remaining 30 % of the samples were allocated to the test data set, distributed as follows: 54 samples for the training data set, 12 NHOSP, 21 HOSP, 7 ICU and 14 EXI, and 22 samples for the test data set 4 samples from NHOSP, 9 HOSP, 3 ICU and 6 EXI. The partitioning of the data was done by computing the randomisation of the samples and ensuring that the randomisation results were reproducible after setting a seed in the programming environment.

To optimize the performance of the classifier, three different machine learning pipelines were initially explored that integrate multiple classification algorithms, and different combinations of features and cross-validation methods. The best results were obtained with the sequential combination of two classification methods: LDA and SVM (Supplementary Figure 11). The total number of classifiers during the optimisation of a combined classifier, considering all proteins, was 253. The goal was that the accuracy associated with the cross-validation data set should be higher than that of the test data set and that the accuracy should be relatively high in both, indicating that there was no overfitting in any data subset. The resulting protein combination was: A2MG, RET4, CYTC, CO5, CLUS, FETUA, FIBG, FIBB, GNPTG, FINC, K1C16, ALBU, K1C9, and IBP2 (Supplementary Figure 12). Starting from this protein panel, the next step allowed the improvement of the classification accuracy of the SVM model. This was achieved by optimizing a combination of the hyperparameters C (0.31) and sigma (0.1601) through a holistic random search. Overall, this is a heuristic method as testing all possible combinations of proteins and hyperparameters would not be computationally feasible. The area under the ROC curve was 0.7355 (Figure 7A) an acceptable performance, considering the challenging classification of the hospitalized group that includes patients of very different severity, as can be assessed in the confusion matrices (Figure 7B). The three LDA dimensions of the model contribute asymmetrically to discriminate the different classes, i.e., each dimension (LD1, LD2, or LD3) can improve the classification accuracy of specific classes (Figure 7C). Similarly, the features contribute asymmetrically to the classification accuracy, according to their corresponding importance value (Figure 7D).

**Figure 7.**
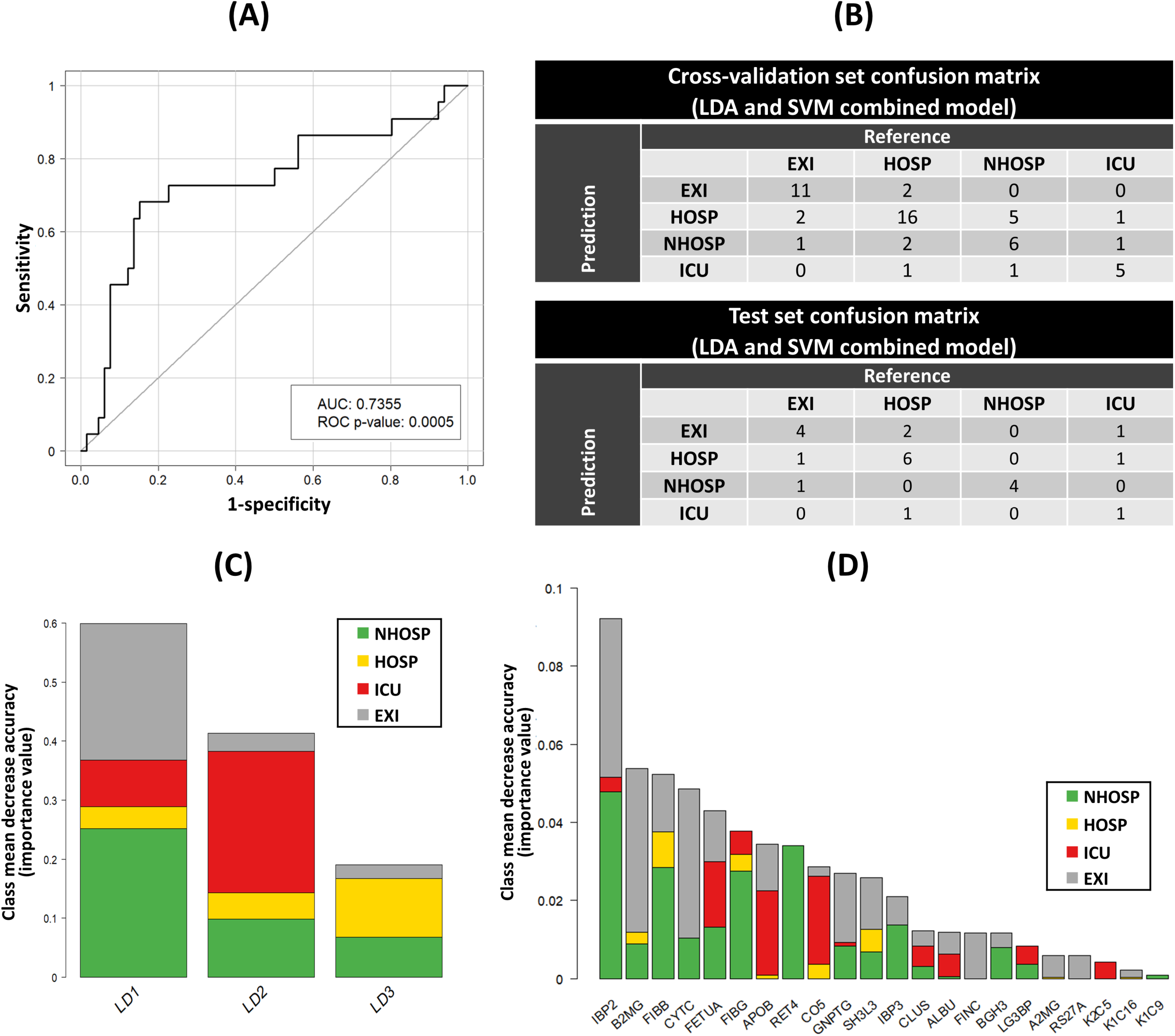
Combined linear discriminant analysis (LDA) and support vector machine (SVM) classifier. (A) ROC curve of the classification efficiency of the combined LDA and SVM model on test set samples. The ROC curves and their associated AUC value and p-value are calculated from the number of samples correctly classified from a data set, taking into account the total number of classes and the number of samples proportionally classified in each group. (B) Confusion matrix of the cross-validation set (top) and test set (bottom) classification by the combined LDA and SVM model. (C) Feature importance bar chart of the random forest LDA dimensions from the combined LDA and SVM classification model. The overall importance of each dimension in classifying the samples is represented as the sum of the random forest mean decrease accuracy measure associated with each class. (D) Importance of all features measured by random forest on the entire dataset. This plot illustrates how proteins contribute to the performance of the classification model. Colour code: green: non-hospitalized (NHOSP); yellow: hospitalised (HOSP); red: Intensive Care Unit (ICU); grey: deceased (EXI).

LDA is a dimensional reduction method that depends on the calculation of the covariance matrices per class, having as final result a set of eigenvalues and eigenvectors (LDA biplots). The eigenvectors are a set of factors, each of which is associated with a feature (i.e., protein), which has a specific weight on the final value of the Linear Discriminant. Therefore, they make possible to represent the performance of the LDA facilitating the interpretation of the model by plotting two-dimensionally each paired combination of Linear Discriminants (Figure 8). This allows the discrimination between the different disease groups, considering a combination of different dimensions. Overall, longitudinal proteomic studies of SARS-CoV-2 infected human cells in culture provided a comprehensive description of the cellular proteome remodeling along the infection. In addition, a panel of differential proteins was identified as candidate biomarkers that allowed the establishment of a method to classify disease severity in a cohort of infected patients, providing new opportunities for their clinical management.

**Figure 8.**
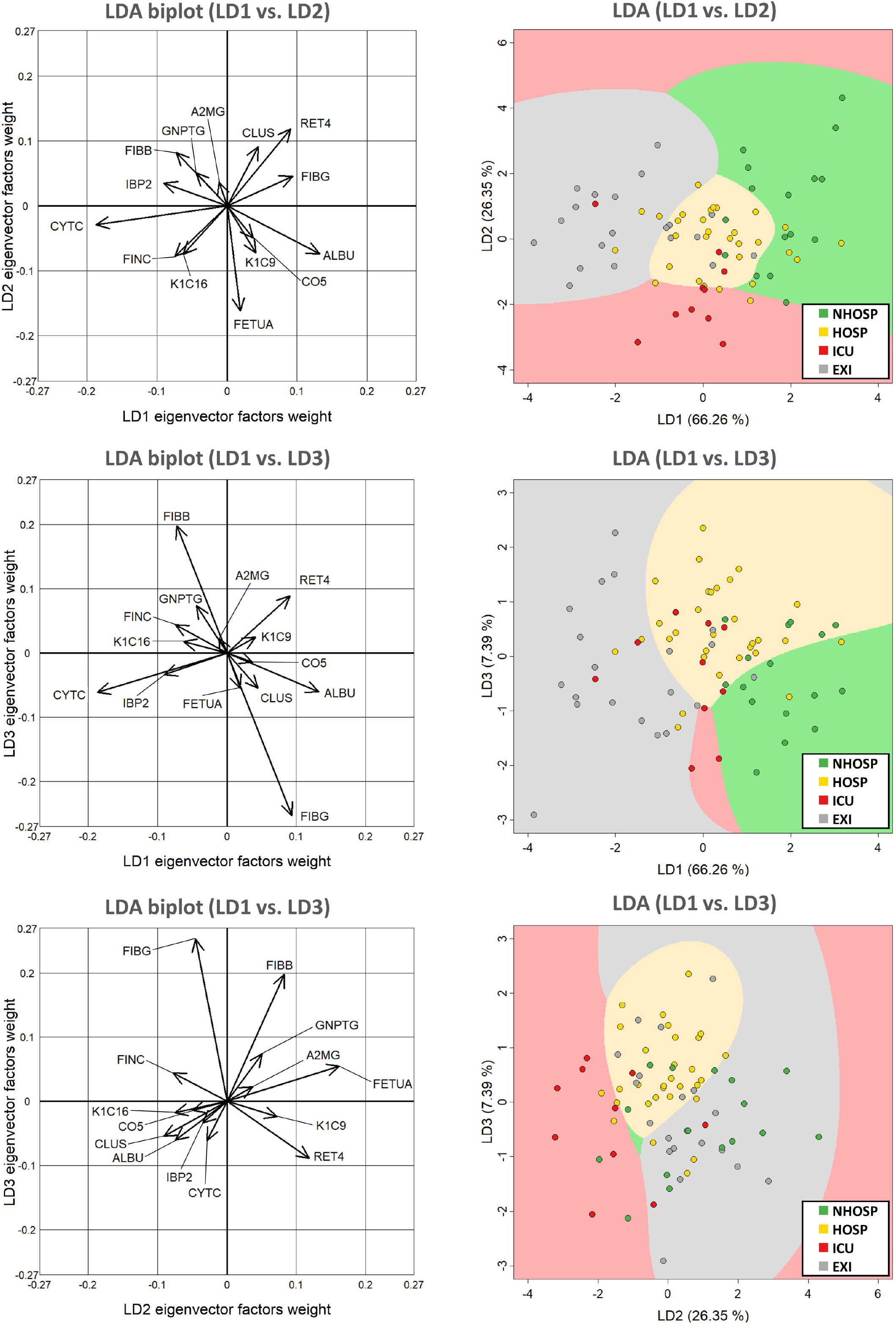
Two-dimensional representation of the results of the combined LDA and SVM classification model. On the left, representations of how the eigenvector factors contribute as proportions to the linear LDA combinations. Given that LDA produces three dimensions or linear discriminants, there are three possible combinations without repetition of two features out of a total of three features. Each of the two-dimensional plots represents one of the combinations of LDA dimensions. The weight associated with each protein varies between the eigenvectors of each linear discriminant. Right, two-dimensional plots of each combination of linear discriminants corresponding to the arrow plots on their left. Samples are represented as points and the colour of the point indicates the corresponding class. The coloured area is the space detected by the SVM model where a sample is more likely to belong to a particular class by its coordinates in the LDA. The percentage of variance of the data retained by each linear discriminant after training the LDA model is shown in the axis names between brackets. Colour code: green: non-hospitalised (NHOSP); yellow: hospitalised (HOSP); red: Intensive Care Unit (ICU); grey: deceased (EXI).

## Discussion

In this study, we aimed to identify proteins regulated by SARS-CoV-2 in human lung epithelial cells that are associated with cellular processes driving the viral infection cycle. Based on the defined mechanistic background, we selected a panel of relevant proteins during infection that can be detected in plasma, using machine learning tools. These proteins were tested as severity markers of COVID-19 in serum samples from a cohort previously characterized in our laboratory (5). The combination of functional studies to understand the host cell response with statistical estimation to hypothesize potential biomarkers is emerging as a promising strategy to define robust disease-associated proteins with potential clinical applications (27).

To fully understand a complex biological process, it is essential to unravel the dynamics of the molecular events involved in order to deduce the hierarchical interplay between them. We describe here a deep time-dependent proteome rewiring that drives a sequential reprogramming of cell biology to allow the progress of SARS-CoV-2 infection, which is in very good agreement with previous reports. Evasion of the cellular innate immunity, control of the translation machinery, regulation of the endocytic and secretory pathways, control of the apoptotic response and autophagy, induction ER stress and UPR, interference with the control of cell polarity and epithelial cell-cell junction integrity, lipid metabolism are the masterpieces of the host cell reprogramming by the virus, that is mediated by key viral proteins (22, 28–36). In addition to the expected accumulation of viral proteins over time, we also detected a time-dependent phosphorylation increase regulatory impact of which is unclear. Phosphorylation, as well as other posttranslational modifications, might influence the protein-protein interaction landscape of viral proteins, playing an essential role in coordinating the virus-cell interaction. As an example, it has been demonstrated that the phosphorylation of nucleoprotein at S79 promotes the interaction of PIN1, regulating positively the production of viral particles or at S51 modulating the binding of RNA to the NTD domain (37). Among the phosphorylation sites previously identified in the nucleoprotein (Supplementary Figure 4), there is no previous reference, to the best of our knowledge, to the S210 and S216 phosphosites described here. These residues are located at a highly disordered region in the C-terminal domain of the protein that is neither involved in RNA binding nor in dimerization and, therefore, the functional implications of these modifications will need additional studies.

The identification of circulating biomarkers to diagnose and predict the severity of COVID-19 and long-term complications has been a priority for the scientific community since the breakthrough of the SARS-CoV-2 pandemic. Different proteome-wide studies have been conducted in this endeavor on different biological fluids including gargle, saliva, CSF, and serum samples (38, 39) (reviewed in 5). Although systematic studies involving larger cohorts are needed, several differential proteins have been commonly reported and, therefore, might be considered as potential biomarkers (Supplementary table 7). Most of these candidates emerge from the statistical assessment of heterogeneous cohorts and might reflect a systemic condition, as biological fluids collect information from the whole organism. Proteome analyses of target cells exposed to SARS-CoV-2 provide a functional framework for the selection of infection-associated proteins and therefore, enhance their specificity as biomarkers whose discriminatory capacity could be then validated in patient samples accessible by minimally invasive methods. In the last few years, machine learning strategies have been increasingly employed for unbiased feature selection from proteome-wide datasets, in particular for SARS-CoV-2 studies (40, 41). In this study, we selected 30 differential proteins in A549-ACE2 cells exposed to SARS-CoV-2, which have been previously detected in plasma/serum according to PAXdb. The selection process was done by recursive random forest based on the accumulated importance value of variables after 50 iterations. PRM analysis allowed the quantification of 22 of these proteins in serum samples from patients with increasing COVID-19 severity that was previously studied in our laboratory (5).

Starting from the PRM-derived protein abundance data, LDA and SVM were sequentially used to optimize a classifier to stratify COVID-19 serum patients according to the severity of the disease. LDA classification provided higher accuracy associated with the cross-validation and test set simultaneously than other classifiers tested in this analysis. Therefore, LDA was applied as a dimensional reduction technique and feature extraction method to select relevant proteins that are used in a second classification algorithm, SVM, which uses a radial kernel. Though the overall important value decreased from LDA1 to 3, each specific LDA is particularly important for classifying samples belonging to certain classes. The final model included 14 proteins with an area under the ROC curve of 0.7355 whose association with COVID-19 progression can be postulated based on their differential expression in A549-ACE2 cells exposed to SARS-CoV-2. Of the 14 proteins, some (A2GM, RET4, CYTC, FETUA, IBP2, FIBB, FIBG, FINC, CO5, and ALBU) have already been proposed as indicators of COVID-19 severity and associated comorbidities (5). Previous studies have demonstrated that acute renal failure is a common complication of COVID-19, affecting up to 46% (42). The increase of A2GM and the concomitant decrease of ALBU suggest the progression of renal damage, a hypothesis that is further supported by the up-regulation of CYTC and IBP2 (43). From the early stages of the pandemic, coagulation abnormalities were recognized as one of the complications of COVID-19, with an estimate of thrombotic events in up to 60% of the patients, with the highest incidence in those individuals with severe disease and requiring an ICU admission (44, 45). According to this evidence, the observed accumulation of FIBB and FIBG may serve as a measure of coagulation changes associated with the progression and severity of COVID-19. To the best of our knowledge, the additional 4 proteins of the classifier (CLUS, K1C16, K1C9, and GNPTG) have not been associated so far with COVID-19 progression. Keratin intermediate filaments are essential elements of epithelial cells and tissue architecture. Downregulation of K1C16 and K1C9 in A549-ACE2 cells during SRS-CoV-2 infection, as well as the severity-dependent drop observed in the serum of COVID-19 patients, might be associated with the lung epithelial dysfunction resulting from the inflammatory reaction that leads to the acute respiratory distress syndrome (46). The progressive decrease of these proteins in serum might also explain the skin manifestations that have been observed in some COVID-19 patients (47). CO5 up-regulation is likely part of the well-known activation of the complement cascade in COVID-19 (48) as well as the decrease of CLUS. CLUS is an extracellular chaperone that can alter the membrane attack complex (MAC) formation preventing complement-mediated cytolysis (49). Finally, as for other coronaviruses, SARS-CoV-2 virion assembly occurs into the ER-Golgi intermediate compartment (ERGIC) (50, 51). However, it has been recently shown that, differently from other enveloped viruses, SARS-CoV-2 and other betacoronaviruses follow a non-lytic lysosomal exocytotic pathway once assembled in the ERGIC compartment (52). GNPTG plays an essential role in the trafficking of proteins from the Golgi to the lysosome (53) and, therefore, it is tempting to speculate that it may play a central role in the assembly and exocytosis of SARS-CoV-2 viral particles.

Overall, we have identified proteins and phosphoproteins in A549-ACE2 lung epithelial cells that are regulated by SARS-CoV-2 and provide molecular evidence to understand the host cell response. A panel of the differentially regulated proteins that are measurable in blood was verified in human serum samples from COVID-19 patients by PRM-targeted mass spectrometry. Changes in the abundance of 22 proteins parallel our findings in the cellular model, indicating their association with the host response to SARS-CoV-2 infection. PRM data were then analyzed with machine learning tools resulting in an optimized classifier of 14 proteins to stratify patients according to their severity. Remarkably, these proteins summarize some of the well-known processes associated with SARS-CoV-2 infection.

## Materials and methods

### Cells

Human lung adenocarcinoma cells (A549) expressing human ACE2 and Vero-E6 cells have been previously described. Cells were maintained complete media (Dulbecco-s Modified Eaglés medium; DMEM) supplemented with 10 mM HEPES, 1X non-essential amino acids (GIBCO), 100 U/mL penicillin-streptomycin (GIBCO) and 10% fetal bovine serum (FBS; heat-inactivated at 56°C for 30 min). Unless otherwise stated, all infection experiments were performed at 37°C in a CO2 incubator (5% CO2) the presence of 2% FBS in the absence of selection antibiotics.

### Viruses

SARS-CoV-2 (Coronaviridae; Orthocoronavirinae; Betacoronavirus; Sarbecovirus; strain NL/2020) was provided by Dr. R. Molenkamp, Erasmus University Medical Center Rotterdam. SARS-CoV-2 stocks were produced in Vero-E6 cells by inoculation at a multiplicity of infection (MOI) of 0.001 TCID50/cell. Cell supernatants were harvested at 48 hpi, cleared by centrifugation, aliquoted, and stored at −80°C. SARS-CoV-2 virus titers were determined by endpoint dilutions and immunofluorescence microscopy using an antibody against N protein (nucleoprotein), as previously described (54).

### Single Cycle Infection experiments

A549-ACE2 cells were inoculated at a multiplicity of infection of 1FFU/cell (MOI 1) by incubation with infectious supernatants for 1 hour at 37°C. Inoculum was removed and cells were washed with warm PBS before replenishing the cells with complete media containing 2% FBS. Samples of the cells were collected at the indicated time points by scraping the cells in ice-cold 1X PBS after media removal and washing with ice-cold PBS. Cell pellets were produced by centrifugation at 4500 xg/5 min/4°C. Cell pellets were resuspended in lysis buffer (see below). All infection procedures were carried out following international guidelines in an authorized BSL3 facility under the supervision of the local Biosafety Committee. All activities were authorized by CSIC Ethics Committee and the Comisión Interministerial de Bioseguridad from the Spanish Government.

### Cell sample preparation

A549-ACE2 cells (2 × 10^6^ cells per biological replicate) were lysed with 5% sodium dodecyl sulfate (SDS) and 25 mM triethylammonium bicarbonate (TEAB) supplemented with 10 mM tris(2-carboxyethyl)phosphine (TCEP) and 10 mM chloroacetamide (CAA)). The samples were incubated at 60°C for 1h, followed by sonication using an ultrasonic processor UP50H (Hielscher Ultrasonics) for 1 min on ice (0.5 cycles, 100% amplitude). The protein extracts were centrifuged at 18,400 × g for 10 min and the supernatant was transferred to a new tube and quantified by PIERCE 660 nm reagent (Thermo Scientific) supplemented with ionic Detergent Compatibility Reagent (Thermo Scientific). Protein digestion on S-Trap columns (Protifi) was performed following the manufacturer’s instructions with minor changes (55, 56). Briefly, 80 µg of each sample were digested at 37°C overnight using trypsin:protein ratio of 1:15. After digestion, tryptic peptides were quantified by fluorimetry using QuBit (Thermo Fisher Scientific), according to the manufacturer’s instructions, and 27 µg were labeled using Tandem Mass Tags (TMT)pro™ 16plex kit. For this purpose, tryptic peptides of each sample were resuspended with 25 mM TEAB buffer and 50% anhydrous acetonitrile (ACN) and labeled with distinct TMT label reagents, (Supplementary Figure 1). A pool of all digested samples (1.8 µg/each sample) was labeled with 134N to be included in the experiment as an internal standard (IS). After 2h, TMT reaction was quenched with 0.3% hydroxylamine for 15 minutes at room temperature and, after that, the individual samples were combined in equal parts. For phosphopeptides enrichment, 270 µg were transferred to a new tube, dried in a speed vacuum, and frozen until further processing. In turn, 80 µg were kept for the fractionation using styrene Divinylbenzene reverse phase sulfonate (SDB-RPS) STAGE Tips.

### Basic-pH fractionation using SDB-RPS STAGE Tips

The fractionation of TMT-labeled peptides was performed using in-home-made STAGE tips prepared from SDB-RPS solid-phase extraction disks (Empore™) as previously reported (57). SDB-RPS solid-phase extraction disks were packed into a 200 µL tip according to the STAGE tip protocol described by our research group (56). The STAGE tip was inserted onto the top of a 2-mL tube using an in-home-made adapter, activated with 100 µL MeOH, and centrifuged at 900 × g for 3 min. The tip was then conditioned with steps of 100 µL 50% ACN and 0.1% formic acid (FA) and centrifuge, followed by three equilibration steps with 0.1% FA. After that, TMT-labeled peptides (80 µg) were reconstituted in 100 µL 1% FA (pH < 3) and loaded onto the STAGE tip. The sample was centrifuged at 900 × g for 5 min and the collected flow-through was loaded again to improve the peptide recovery yield. The STAGE tip was washed with 100 µL 0.1% FA, followed by 100 µL H2O. The elution was carried out using a 10-stepwise elution with 100 µl of 5 mM ammonium formate buffer and increasing acetonitrile concentrations (0, 5.0%, 7.5%, 10.0%, 12.5%, 15.0%, 17.5%, 20%, 25%, and 45%). Fractions were dried in a speed vacuum and frozen until further processing.

### TiO2 phosphopeptide enrichment

Phosphopeptide enrichment was carried out as described previously (58). The TiO2 slurry was prepared by mixing 10 mg Titansphere beads (10 µm, GL Sciences) with 80 µl of 1M glycolic acid in 80% acetonitrile and 1% Trifluoroacetic acid (TFA) for 30 min at 25oC. After being resuspended with 600 µl 60% ACN and 1% TFA, 250 µg TMT-labeled peptides were incubated at room temperature with a volume of TiO2 slurry corresponding to a TiO2:protein ratio of 10:1. Following a 25-minute incubation period, the beads were spined down in a bench-top centrifuge and the flow-through was removed (non-phosphorylated peptides). The phosphopeptides bound to the titansphere beads were resuspended in 150 µl of 60% ACN and 1% TFA and transferred to a 200 µL tip with a 10 µm filter (MoBiTec). The wash step was repeated and the phosphopeptides were subsequently eluted using 5% ammonium hydroxide (NH4OH) (2x) and 25% ACN in 10% NH4OH (2x). The eluted fractions were pooled and acidified with TFA. Phosphopeptide enriched fraction was subsequently desalted using an in-house packed Oligo R3 reversed-phase micro-column, dried and stored until further processing.

### Analysis by liquid chromatography coupled to mass spectrometry

Each sample (10 fractions or phosphopeptide enriched-fraction) was quantified by fluorimetry (QuBit) and 1 µg was individually analyzed by nano-Liquid Chromatography coupled to Electrospray Ionization Tandem Mass Spectrometry (nanoLC-ESI-MS/MS) analysis using an Ultimate 3000 nano HPLC system (Thermo Fisher Scientific) coupled online to an Orbitrap Exploris™ 240 mass spectrometer (Thermo Fisher Scientific). The fractions (1 µg in 5 µl of injection volume) were loaded on a 50 cm × 75 μm Easy-spray PepMap C18 analytical column at 45°C and were separated at a flow rate of 300 nL/min using a 120 min gradient ranging from 2 % to 95 % mobile phase B (mobile phase A: 0.1% formic acid (FA); mobile phase B: 80 % acetonitrile (ACN) in 0.1% FA). To avoid carry-over, two 40 min blank samples (mobile phase A) were systematically run between samples. Data acquisition was performed using a data-dependent top method, in full scan positive mode, scanning 375 to 1200 m/z. MS1scans were acquired at an orbitrap resolution of 60,000 at m/z 200, with a normalized automatic gain control (AGC) target of 300%, a radio frequency (RF) lens of 80%, and an automatic maximum injection time (IT). The top 20 most intense ions from each MS1 scan were selected and fragmented with a Higher-energy collisional dissociation (HCD) of 30%. Resolution for HCD spectra was set to 45,000 at m/z 200, with an AGC target of 100% and an automatic maximum IT. Isolation of precursors was performed with an isolation window of 0.7 m/z and 45 s of exclusion duration. Precursor ions with single, unassigned, or six and higher charge states from fragmentation selection were excluded. For phosphopeptide analysis, HCD was set to 32% and the exclusion duration was decreased to 30 s.

### Shotgun data analysis

Data obtained by mass spectrometry were analyzed with Proteome Discoverer (v2.5.0.400) using four search engines (Mascot (v2.7.0), MsAmanda (v2.4.0), MsFragger (v3.1.1), and Sequest HT) using a target/decoy *Homo sapiens* + *SARS-CoV2* Uniprot Knowledgebase database (25th February 2021, 20,462 sequences) with the most common laboratory contaminants (cRAP database with 69 sequences). Search parameters were set as follows: cysteine carbamidomethyl (+57.021464 Da) and TMT6plex (+229.162932 Da) on lysine and N-term as fixed modifications; methionine oxidation (M) (+15.994915 Da), N-term acetylation (+42.010565 Da), and Gln→pyro-Glu (-17.026549 Da) as variable modifications. Precursor and fragment mass tolerances were set at 10 ppm and 0.02 Da respectively, and trypsin/P was selected as a protease with a maximum of 2 missed cleavage sites. The false discovery rate (FDR) for proteins, peptides, and peptide spectral matches (PSMs) peptides was kept at 1%. The quantitation was also performed in Proteome Discoverer using the “Reporter Ions Quantifier” feature in the quantification workflow using the following parameters: unique+razor peptides were used for quantitation, co-isolation threshold was set at 50%, signal to noise of reporter ions was 10, and the normalization and scaling were performed considering the total peptide amount and the control (IS) average, respectively. The protein ratio was calculated considering the protein abundance and the hypothesis test was based on a t-test (background-based). Protein groups (master proteins) with an FDR lower than 1% and with abundance values in both IS were considered for quantitation. A p-value≤0.05 adjusted using Benjamini-Hochberg was set to determine the proteins found differentially expressed at 3h, 6h, 9h, and 16h compared with the Mock-infected cells. Volcano plot and Principal Component Analysis (PCA) were performed in Proteome Discover considering the differentially expressed proteins for each comparison. Functional analysis of differentially expressed and phosphorylated proteins (peptides) was performed using Metascape and Coronascape (59). Transcription Factor Enrichment Analysis (TFEA) and Kinase Enrichment Analysis (KEA) were performed using the Expression2Kinases webtool (X2Kweb; https://maayanlab.cloud/X2K/) (60, 61).

### Feature selection for patient stratification

Proteins whose presence in plasma has been already reported were retrieved from the total list of identified proteins according to the abundance information from paxdb (URL: https://pax-db.org/dataset/9606/1394854118/) (62). A classifier was then generated with random forest considering the 48 proteins to classify the different cell groups exposed to SARS-CoV-2, according to the following pipeline. Intensity data were normalised by Proteome Discoverer™ and then, log2 was calculated. All data were fed into a recursive random forest algorithm to rank the proteins by importance. Random forest was run for 30 rounds to eliminate iteratively those proteins with the lower important value, until only one protein was retained. Then, the dataset was regenerated and the whole process of iterative variable elimination started again up to 50 times, accumulating the importance value from each iteration. Proteins were ranked according to the accumulated importance and the top 30 proteins were selected for further experimental testing in human samples (Supplementary Table 5).

### Serum processing using automatic SP3-Based Protein Digestion

Serum samples were diluted with stock buffer to a final concentration of 2.5% SDS, 25 mM Triethylammonium bicarbonate (TEAB), 5 mM TCEP, and 10 mM chloroacetamide (CAA) and aliquoted into 8-well strips. Samples were incubated at 60oC for 30 min in a Jitterbug 4 incubator (Fisher Scientific) for protein reduction and alkylation. For automatic SP3-based protein digestion, 50 µg of protein was processed in the OT-2 platform as previously reported (63). Briefly, 100 µg of MagReSyn® Amine beads (20 µg/µl suspension in 20% ethanol) were mixed with 30 µl of serum and 70 µl ACN. The protein-bead aggregation was carried out by two steps of 10 min incubation at room temperature without agitation. Protein-bead aggregates were washed three times with 100 µl ACN, followed by three washes with 70% ACN. For protein digestion, samples were incubated for 16h at 37°C with a mix of 1.5 µg Trypsin and 0.10 µg Lys-C (125-05061, WAKO) prepared in 25 mM TEAB. Following digestion, supernatants containing peptides were transferred to a new tube and acidified at a final concentration of 1% (v/v) formic acid. Peptides were dried in a speed vacuum and conserved at -20°C until further processing.

### Optimization of the Parallel Reaction Monitoring (PRM) method

Of the thirty proteins selected by random forest, a list of twenty-nine proteins to be monitored by parallel reaction monitoring method (PRM) was imported into Skyline v.22.2.0.255. The selection of 3 proteotypic peptides per protein was made considering the information of MS/MS spectral libraries in SRM Atlas and previous DDA experiments with plasma/serum according to the following criteria: peptide length between 7 and 25 residues, no missed cleavages, and exclusion of peptides containing methionine, histidine, and other amino acids susceptible to undergoing modifications (except for cysteine). To generate the experimental peptide library within Skyline, a serum pool containing synthetic peptides to be monitored (external standard) was analyzed by 3 PRM unscheduled sub-methods exported from Skyline v.22.2.0.255. For this purpose, 5 µl of sample (corresponding to 1 µg of peptide) were loaded on a 50 cm × 75 μm Easy-spray PepMap C18 analytical column at 45°C using an Ultimate 3000 nano HPLC system (Thermo Fisher Scientific) coupled online to an Orbitrap Exploris™ 240 mass spectrometer (Thermo Fisher Scientific). Peptides were separated at a flow rate of 300 nL/min using a 60 min gradient ranging from 2 % to 95 % mobile phase B (mobile phase A: 0.1% formic acid (FA); mobile phase B: 80 % acetonitrile (ACN) in 0.1% FA). Data acquisition was performed in PRM mode for monitoring the 86 peptides (approximately 29 peptides per method). The selected peptides were fragmented with a Higher-energy collisional dissociation (HCD) of 30%. The isolation window was set to 0.8 and the resolution to 30,000. The obtained data were searched against a target/decoy Homo sapiens+SARS-CoV2 Uniprot Knowledgebase database (25th February 2021, 20,462 sequences) using Mascot (v2.7.0) within Proteome Discoverer (v2.5.0.400). The generated .msf and .pd files were imported into Skyline software for library generation. To optimize the PRM method, serum samples were analyzed using the same procedure to select the peptides that can be experimentally detected and quantified. The raw data was imported into Skyline software. Undetected peptides and proteins (not identified in proteome discoverer) or/and peptides with wide or unquantifiable peaks were removed from the analysis. The final method consisted of a scheduled method of 90 min for monitoring 61 peptides and 23 Proteins (Supplementary Table 5).

### Parallel Reaction Monitoring (PRM) analysis

Tryptic peptides were dried in a speed-vacuum system and resuspended at 200 ng/μl, according to QUBIT quantification (Thermofisher Scientific). Each sample (1 µg of peptides in 5 µl of injection volume) was loaded online on a C18 PepMap 300 µm I.D. 0.3 × 5 mm trapping column (5 µm, 100 Å, Thermo Scientific) and analyzed by nanoLC-ESI-MS/MS analysis using a Thermo Ultimate 3000 RSLC nanoUPLC coupled online to an Orbitrap Exploris™ 240 mass spectrometer (Thermo Fisher Scientific). Peptides were then separated on a 15 cm × 75 μm Easy-spray PepMap C18 analytical column at 45°C with a flow rate of 300 nL/min using a 90 min gradient ranging from 2 % to 95 % mobile phase B (mobile phase A: 0.1% formic acid (FA); mobile phase B: 80 % acetonitrile (ACN) in 0.1% FA). A pool (external standard) was run for each batch of 8 samples to monitor oscillations in the MS signal and in the retention time. The selected peptides (Supplementary Table 2) were fragmented with an HCD of 30%. The isolation window was set to 0.8 and the resolution to 30,000.

### PRM data Analysis

Raw MS data were imported into Skyline and the automatically selected transitions were manually revised considering the intensity distribution of peaks from the fragmentation spectrum contained in the experimental peptide library. Peptides not detected in the most of samples (not identified in proteome discoverer) or/and with wide peaks were removed from the analysis. The peptide area was normalized by dividing the peptide area quantified in each sample by the area of the same peptide in the external standard analyzed in the same batch. The total normalized area of each peptide was calculated by summing all the normalized areas of all associated peptides (each one resulting from the sum of all transitions). Statistical analysis by one-way ANOVA (Tukey’s HSD) & post-hoc tests

### Machine learning modeling of PRM results

All calculations were performed in the R environment, version 4.1.3. All random forest-dependent results were obtained through the “randomForest” library v4.7-1.1. The “caret” library v6.0-93 was used to create the machine learning models. The “MASS” package v7.3-58.1 was used to generate the linear discriminant analysis (LDA) models. The “e1071” library v1.7-13 was used to build the support vector machine (SVM) classifiers. All SVMs created use a radial kernel. The hyperparameters of all classification models were selected by grid search of C and sigma values. The verification package v1.42, was used to create the ROC (receiver operating characteristic) curves. These were generated based on the probability values of the samples assigned to each class as a vector, plus a vector of information about the actual class of each sample. These vectors were entered into the “roc.plot” function to generate the ROC curve.

## Acknowledgments

The CNB was supported by Comunidad de Madrid Grants B2017/BMD-3817 and 2022/BMD-7232. Intramural CSIC PIE/COVID-19 projects 202020E079 and 202020E108. MICIN PID2021-127496NB-100. This research work was also funded by the European Commission—NextGenerationEU (Regulation EU 2020/2094), through CSIC’s Global Health Platform (PTI Salud Global) and Conexión Cancer. We acknowledge R. Molenkamp (Erasmus University Medical Center, Rotterdam, The Netherlands; participant of the EU-funded EVA-GLOBAL project, grant agreement 871029) for the SARS-CoV-2 strain NL/2020 virus.

## References

1. V’kovski P, Kratzel A, Steiner S, Stalder H, Thiel V. 2021. Coronavirus biology and replication: implications for SARS-CoV-2, vol 19, p 155–170. Nature Research.

2. Hamre D, Procknow JJ. 1966. A new virus isolated from the human respiratory tract. Proc Soc Exp Biol Med 121:190–3.

3. McIntosh K, Dees JH, Becker WB, Kapikian AZ, Chanock RM. 1967. Recovery in tracheal organ cultures of novel viruses from patients with respiratory disease. Proc Natl Acad Sci U S A 57:933–40.

4. Struwe W, Emmott E, Bailey M, Sharon M, Sinz A, Corrales FJ, Thalassinos K, Braybrook J, Mills C, Barran P. 2020. The COVID-19 MS Coalition—accelerating diagnostics, prognostics, and treatment, vol 395, p 1761–1762. Lancet Publishing Group.

5. Nunez E, Orera I, Carmona-Rodriguez L, Pano JR, Vazquez J, Corrales FJ. 2022. Mapping the Serum Proteome of COVID-19 Patients; Guidance for Severity Assessment. Biomedicines 10.

6. Völlmy F, van den Toorn H, Chiozzi RZ, Zucchetti O, Papi A, Volta CA, Marracino L, Sega FVD, Fortini F, Demichev V, Tober-Lau P, Campo G, Contoli M, Ralser M, Kurth F, Spadaro S, Rizzo P, Heck AJR. 2021. A serum proteome signature to predict mortality in severe covid-19 patients. Life Science Alliance 4.

7. Demichev V, Tober-Lau P, Lemke O, Nazarenko T, Thibeault C, Whitwell H, Röhl A, Freiwald A, Szyrwiel L, Ludwig D, Correia-Melo C, Aulakh SK, Helbig ET, Stubbemann P, Lippert LJ, Grüning NM, Blyuss O, Vernardis S, White M, Messner CB, Joannidis M, Sonnweber T, Klein SJ, Pizzini A, Wohlfarter Y, Sahanic S, Hilbe R, Schaefer B, Wagner S, Mittermaier M, Machleidt F, Garcia C, Ruwwe-Glösenkamp C, Lingscheid T, Bosquillon de Jarcy L, Stegemann MS, Pfeiffer M, Jürgens L, Denker S, Zickler D, Enghard P, Zelezniak A, Campbell A, Hayward C, Porteous DJ, Marioni RE, Uhrig A, Müller-Redetzky H, Zoller H, Löffler-Ragg J, et al. 2021. A time-resolved proteomic and prognostic map of COVID-19. Cell Systems 12:780–794.e7.

8. Geyer PE, Arend FM, Doll S, Louiset ML, Virreira Winter S, Müller-Reif JB, Torun FM, Weigand M, Eichhorn P, Bruegel M, Strauss MT, Holdt LM, Mann M, Teupser D. 2021. High-resolution serum proteome trajectories in COVID-19 reveal patient-specific seroconversion. EMBO Molecular Medicine 13.

9. Galbraith MD, Kinning KT, Sullivan KD, Baxter R, Araya P, Jordan KR, Russell S, Smith KP, Granrath RE, Shaw JR, Dzieciatkowska M, Ghosh T, Monte AA, D’alessandro A, Hansen KC, Benett TD, Hsieh EWY, Espinosa JM. 2021. Seroconversion stages COVID19 into distinct pathophysiological states. eLife 10.

10. Pruijssers AJ, Denison MR. 2019. Nucleoside analogues for the treatment of coronavirus infections. Curr Opin Virol 35:57–62.

11. Siegel D, Hui HC, Doerffler E, Clarke MO, Chun K, Zhang L, Neville S, Carra E, Lew W, Ross B, Wang Q, Wolfe L, Jordan R, Soloveva V, Knox J, Perry J, Perron M, Stray KM, Barauskas O, Feng JY, Xu Y, Lee G, Rheingold AL, Ray AS, Bannister R, Strickley R, Swaminathan S, Lee WA, Bavari S, Cihlar T, Lo MK, Warren TK, Mackman RL. 2017. Discovery and Synthesis of a Phosphoramidate Prodrug of a Pyrrolo[2,1-f][triazin-4-amino] Adenine C-Nucleoside (GS-5734) for the Treatment of Ebola and Emerging Viruses. J Med Chem 60:1648–1661.

12. Gordon CJ, Tchesnokov EP, Feng JY, Porter DP, Gotte M. 2020. The antiviral compound remdesivir potently inhibits RNA-dependent RNA polymerase from Middle East respiratory syndrome coronavirus. J Biol Chem 295:4773–4779.

13. Sheahan TP, Sims AC, Zhou S, Graham RL, Pruijssers AJ, Agostini ML, Leist SR, Schafer A, Dinnon KH, 3rd, Stevens LJ, Chappell JD, Lu X, Hughes TM, George AS, Hill CS, Montgomery SA, Brown AJ, Bluemling GR, Natchus MG, Saindane M, Kolykhalov AA, Painter G, Harcourt J, Tamin A, Thornburg NJ, Swanstrom R, Denison MR, Baric RS. 2020. An orally bioavailable broad-spectrum antiviral inhibits SARS-CoV-2 in human airway epithelial cell cultures and multiple coronaviruses in mice. Sci Transl Med 12.

14. Gordon CJ, Tchesnokov EP, Woolner E, Perry JK, Feng JY, Porter DP, Gotte M. 2020. Remdesivir is a direct-acting antiviral that inhibits RNA-dependent RNA polymerase from severe acute respiratory syndrome coronavirus 2 with high potency. J Biol Chem 295:6785–6797.

15. Hoffman RL, Kania RS, Brothers MA, Davies JF, Ferre RA, Gajiwala KS, He M, Hogan RJ, Kozminski K, Li LY, Lockner JW, Lou J, Marra MT, Mitchell LJ, Jr., Murray BW, Nieman JA, Noell S, Planken SP, Rowe T, Ryan K, Smith GJ, 3rd, Solowiej JE, Steppan CM, Taggart B. 2020. Discovery of Ketone-Based Covalent Inhibitors of Coronavirus 3CL Proteases for the Potential Therapeutic Treatment of COVID-19. J Med Chem 63:12725–12747.

16. de Vries M, Mohamed AS, Prescott RA, Valero-Jimenez AM, Desvignes L, O’Connor R, Steppan C, Devlin JC, Ivanova E, Herrera A, Schinlever A, Loose P, Ruggles K, Koralov SB, Anderson AS, Binder J, Dittmann M. 2021. A comparative analysis of SARS-CoV-2 antivirals characterizes 3CL(pro) inhibitor PF-00835231 as a potential new treatment for COVID-19. J Virol 95.

17. Tao K, Tzou PL, Nouhin J, Bonilla H, Jagannathan P, Shafer RW. 2021. SARS-CoV-2 Antiviral Therapy. Clin Microbiol Rev 34:e0010921.

18. Hoffmann M, Kleine-Weber H, Schroeder S, Krüger N, Herrler T, Erichsen S, Schiergens TS, Herrler G, Wu NH, Nitsche A, Müller MA, Drosten C, Pöhlmann S. 2020. SARS-CoV-2 Cell Entry Depends on ACE2 and TMPRSS2 and Is Blocked by a Clinically Proven Protease Inhibitor. Cell 181:271–280.e8.

19. Bellino S. 2022. COVID-19 treatments approved in the European Union and clinical recommendations for the management of non-hospitalized and hospitalized patients. Ann Med 54:2856–2860.

20. Wang Q, Zhang Y, Wu L, Niu S, Song C, Zhang Z, Lu G, Qiao C, Hu Y, Yuen KY, Wang Q, Zhou H, Yan J, Qi J. 2020. Structural and Functional Basis of SARS-CoV-2 Entry by Using Human ACE2. Cell 181:894–904 e9.

21. Yan R, Zhang Y, Li Y, Xia L, Guo Y, Zhou Q. 2020. Structural basis for the recognition of SARS-CoV-2 by full-length human ACE2. Science 367:1444–1448.

22. Pizzato M, Baraldi C, Boscato Sopetto G, Finozzi D, Gentile C, Gentile MD, Marconi R, Paladino D, Raoss A, Riedmiller I, Ur Rehman H, Santini A, Succetti V, Volpini L. 2022. SARS-CoV-2 and the Host Cell: A Tale of Interactions. Frontiers in Virology 1.

23. Bojkova D, Klann K, Koch B, Widera M, Krause D, Ciesek S, Cinatl J, Münch C. 2020. Proteomics of SARS-CoV-2-infected host cells reveals therapy targets. Nature 583:469–472.

24. Bouhaddou M, Memon D, Meyer B, White KM, Rezelj VV, Correa Marrero M, Polacco BJ, Melnyk JE, Ulferts S, Kaake RM, Batra J, Richards AL, Stevenson E, Gordon DE, Rojc A, Obernier K, Fabius JM, Soucheray M, Miorin L, Moreno E, Koh C, Tran QD, Hardy A, Robinot R, Vallet T, Nilsson-Payant BE, Hernandez-Armenta C, Dunham A, Weigang S, Knerr J, Modak M, Quintero D, Zhou Y, Dugourd A, Valdeolivas A, Patil T, Li Q, Hüttenhain R, Cakir M, Muralidharan M, Kim M, Jang G, Tutuncuoglu B, Hiatt J, Guo JZ, Xu J, Bouhaddou S, Mathy CJP, Gaulton A, Manners EJ, et al. 2020. The Global Phosphorylation Landscape of SARS-CoV-2 Infection. Cell 182:685–712.e19.

25. Stukalov A, Girault V, Grass V, Karayel O, Bergant V, Urban C, Haas DA, Huang Y, Oubraham L, Wang A, Hamad MS, Piras A, Hansen FM, Tanzer MC, Paron I, Zinzula L, Engleitner T, Reinecke M, Lavacca TM, Ehmann R, Wolfel R, Jores J, Kuster B, Protzer U, Rad R, Ziebuhr J, Thiel V, Scaturro P, Mann M, Pichlmair A. 2021. Multilevel proteomics reveals host perturbations by SARS-CoV-2 and SARS-CoV. Nature 594:246–252.

26. Fritch EJ, Mordant AL, Gilbert TSK, Wells CI, Yang X, Barker NK, Madden EA, Dinnon KH, 3rd, Hou YJ, Tse LV, Castillo IN, Sims AC, Moorman NJ, Lakshmanane P, Willson TM, Herring LE, Graves LM, Baric RS. 2023. Investigation of the Host Kinome Response to Coronavirus Infection Reveals PI3K/mTOR Inhibitors as Betacoronavirus Antivirals. J Proteome Res 22:3159–3177.

27. Babacic H, Christ W, Araujo JE, Mermelekas G, Sharma N, Tynell J, Garcia M, Varnaite R, Asgeirsson H, Glans H, Lehtio J, Gredmark-Russ S, Klingstrom J, Pernemalm M. 2023. Comprehensive proteomics and meta-analysis of COVID-19 host response. Nat Commun 14:5921.

28. Stewart H, Lu Y, O’Keefe S, Valpadashi A, Cruz-Zaragoza LD, Michel HA, Nguyen SK, Carnell GW, Lukhovitskaya N, Milligan R, Adewusi Y, Jungreis I, Lulla V, Matthews DA, High S, Rehling P, Emmott E, Heeney JL, Davidson AD, Edgar JR, Smith GL, Firth AE. 2023. The SARS-CoV-2 protein ORF3c is a mitochondrial modulator of innate immunity. iScience 26:108080.

29. Lenhard S, Gerlich S, Khan A, Rodl S, Bokenkamp JE, Peker E, Zarges C, Faust J, Storchova Z, Raschle M, Riemer J, Herrmann JM. 2023. The Orf9b protein of SARS-CoV-2 modulates mitochondrial protein biogenesis. J Cell Biol 222.

30. Jiao S, Miranda P, Li Y, Maric D, Holmgren M. 2023. Some aspects of the life of SARS-CoV-2 ORF3a protein in mammalian cells. Heliyon 9:e18754.

31. Lopez-Ayllon BD, de Lucas-Rius A, Mendoza-Garcia L, Garcia-Garcia T, Fernandez-Rodriguez R, Suarez-Cardenas JM, Santos FM, Corrales F, Redondo N, Pedrucci F, Zaldivar-Lopez S, Jimenez-Marin A, Garrido JJ, Montoya M. 2023. SARS-CoV-2 accessory proteins involvement in inflammatory and profibrotic processes through IL11 signaling. Front Immunol 14:1220306.

32. Burnap SA, Ortega-Prieto AM, Jimenez-Guardeno JM, Ali H, Takov K, Fish M, Shankar-Hari M, Giacca M, Malim MH, Mayr M. 2023. Cross-Linking Mass Spectrometry Uncovers Interactions Between High-Density Lipoproteins and the SARS-CoV-2 Spike Glycoprotein. Mol Cell Proteomics 22:100600.

33. Min YQ, Huang M, Feng K, Jia Y, Sun X, Ning YJ. 2023. A New Cellular Interactome of SARS-CoV-2 Nucleocapsid Protein and Its Biological Implications. Mol Cell Proteomics 22:100579.

34. Davies JP, Sivadas A, Keller KR, Roman BK, Wojcikiewicz RJH, Plate L. 2024. Expression of SARS-CoV-2 Nonstructural Proteins 3 and 4 Can Tune the Unfolded Protein Response in Cell Culture. J Proteome Res 23:356–367.

35. Zhou Y, Liu Y, Gupta S, Paramo MI, Hou Y, Mao C, Luo Y, Judd J, Wierbowski S, Bertolotti M, Nerkar M, Jehi L, Drayman N, Nicolaescu V, Gula H, Tay S, Randall G, Wang P, Lis JT, Feschotte C, Erzurum SC, Cheng F, Yu H. 2023. A comprehensive SARS-CoV-2-human protein-protein interactome reveals COVID-19 pathobiology and potential host therapeutic targets. Nat Biotechnol 41:128–139.

36. Kim DK, Weller B, Lin CW, Sheykhkarimli D, Knapp JJ, Dugied G, Zanzoni A, Pons C, Tofaute MJ, Maseko SB, Spirohn K, Laval F, Lambourne L, Kishore N, Rayhan A, Sauer M, Young V, Halder H, la Rosa NM, Pogoutse O, Strobel A, Schwehn P, Li R, Rothballer ST, Altmann M, Cassonnet P, Cote AG, Vergara LE, Hazelwood I, Liu BB, Nguyen M, Pandiarajan R, Dohai B, Coloma PAR, Poirson J, Giuliana P, Willems L, Taipale M, Jacob Y, Hao T, Hill DE, Brun C, Twizere JC, Krappmann D, Heinig M, Falter C, Aloy P, Demeret C, Vidal M, Calderwood MA, et al. 2023. A proteome-scale map of the SARS-CoV-2-human contactome. Nat Biotechnol 41:140–149.

37. Wu J, Zhong Y, Liu X, Lu X, Zeng W, Wu C, Xing F, Cao L, Zheng F, Hou P, Peng H, Li C, Guo D. 2022. A novel phosphorylation site in SARS-CoV-2 nucleocapsid regulates its RNA-binding capacity and phase separation in host cells. J Mol Cell Biol 14.

38. Ihling C, Tanzler D, Hagemann S, Kehlen A, Huttelmaier S, Arlt C, Sinz A. 2020. Mass Spectrometric Identification of SARS-CoV-2 Proteins from Gargle Solution Samples of COVID-19 Patients. J Proteome Res 19:4389–4392.

39. Azzi L, Carcano G, Gianfagna F, Grossi P, Gasperina DD, Genoni A, Fasano M, Sessa F, Tettamanti L, Carinci F, Maurino V, Rossi A, Tagliabue A, Baj A. 2020. Saliva is a reliable tool to detect SARS-CoV-2. The Journal of infection 81:e45–e50.

40. Sun H, Li J, Murphy RF. 2024. Expanding the coverage of spatial proteomics: a machine learning approach. Bioinformatics 40.

41. Zhang Y, Bharathi V, Dokoshi T, de Anda J, Ursery LT, Kulkarni NN, Nakamura Y, Chen J, Luo EWC, Wang L, Xu H, Coady A, Zurich R, Lee MW, Matsui T, Lee H, Chan LC, Schepmoes AA, Lipton MS, Zhao R, Adkins JN, Clair GC, Thurlow LR, Schisler JC, Wolfgang MC, Hagan RS, Yeaman MR, Weiss TM, Chen X, Li MMH, Nizet V, Antoniak S, Mackman N, Gallo RL, Wong GCL. 2024. Viral afterlife: SARS-CoV-2 as a reservoir of immunomimetic peptides that reassemble into proinflammatory supramolecular complexes. Proc Natl Acad Sci U S A 121:e2300644120.

42. Chan L, Chaudhary K, Saha A, Chauhan K, Vaid A, Zhao S, Paranjpe I, Somani S, Richter F, Miotto R, Lala A, Kia A, Timsina P, Li L, Freeman R, Chen R, Narula J, Just AC, Horowitz C, Fayad Z, Cordon-Cardo C, Schadt E, Levin MA, Reich DL, Fuster V, Murphy B, He JC, Charney AW, Bottinger EP, Glicksberg BS, Coca SG, Nadkarni GN, on behalf of the Mount Sinai CIC. 2021. AKI in Hospitalized Patients with COVID-19. J Am Soc Nephrol 32:151–160.

43. Mester P, Rath U, Schmid S, Amend P, Keller D, Krautbauer S, Bondarenko S, Muller M, Buechler C, Pavel V. 2024. Serum Insulin-like Growth Factor-Binding Protein-2 as a Prognostic Factor for COVID-19 Severity. Biomedicines 12.

44. Dutch C, Thrombosis C, Kaptein FHJ, Stals MAM, Grootenboers M, Braken SJE, Burggraaf JLI, van Bussel BCT, Cannegieter SC, Ten Cate H, Endeman H, Gommers D, van Guldener C, de Jonge E, Juffermans NP, Kant KM, Kevenaar ME, Koster S, Kroft LJM, Kruip M, Leentjens J, Marechal C, Soei YL, Tjepkema L, Visser C, Klok FA, Huisman MV. 2021. Incidence of thrombotic complications and overall survival in hospitalized patients with COVID-19 in the second and first wave. Thromb Res 199:143–148.

45. Jenner WJ, Kanji R, Mirsadraee S, Gue YX, Price S, Prasad S, Gorog DA. 2021. Thrombotic complications in 2928 patients with COVID-19 treated in intensive care: a systematic review. J Thromb Thrombolysis 51:595–607.

46. Swenson KE, Swenson ER. 2021. Pathophysiology of Acute Respiratory Distress Syndrome and COVID-19 Lung Injury. Crit Care Clin 37:749–776.

47. Mahdiabadi S, Rajabi F, Tavakolpour S, Rezaei N. 2022. Immunological aspects of COVID-19-related skin manifestations: Revisiting pathogenic mechanism in the light of new evidence. Dermatol Ther 35:e15758.

48. Perico L, Benigni A, Casiraghi F, Ng LFP, Renia L, Remuzzi G. 2021. Immunity, endothelial injury and complement-induced coagulopathy in COVID-19. Nat Rev Nephrol 17:46–64.

49. Fox CR, Parks GD. 2021. Complement Inhibitors Vitronectin and Clusterin Are Recruited from Human Serum to the Surface of Coronavirus OC43-Infected Lung Cells through Antibody-Dependent Mechanisms. Viruses 14.

50. de Haan CA, Rottier PJ. 2005. Molecular interactions in the assembly of coronaviruses. Adv Virus Res 64:165–230.

51. Siu YL, Teoh KT, Lo J, Chan CM, Kien F, Escriou N, Tsao SW, Nicholls JM, Altmeyer R, Peiris JS, Bruzzone R, Nal B. 2008. The M, E, and N structural proteins of the severe acute respiratory syndrome coronavirus are required for efficient assembly, trafficking, and release of virus-like particles. J Virol 82:11318–30.

52. Ghosh S, Dellibovi-Ragheb TA, Kerviel A, Pak E, Qiu Q, Fisher M, Takvorian PM, Bleck C, Hsu VW, Fehr AR, Perlman S, Achar SR, Straus MR, Whittaker GR, de Haan CAM, Kehrl J, Altan-Bonnet G, Altan-Bonnet N. 2020. beta-Coronaviruses Use Lysosomes for Egress Instead of the Biosynthetic Secretory Pathway. Cell 183:1520–1535 e14.

53. Khan SA, Tomatsu SC. 2020. Mucolipidoses Overview: Past, Present, and Future. Int J Mol Sci 21.

54. Castro V, Perez-Berna AJ, Calvo G, Pereiro E, Gastaminza P. 2023. Three-Dimensional Remodeling of SARS-CoV2-Infected Cells Revealed by Cryogenic Soft X-ray Tomography. ACS Nano 17:22708–22721.

55. Ciordia S, Alvarez-Sola G, Rullan M, Urman JM, Avila MA, Corrales FJ. 2021. Digging deeper into bile proteome. J Proteomics 230:103984.

56. Ciordia S, Alvarez-Sola G, Rullan M, Urman JM, Avila MA, Corrales FJ. 2022. Bile Processing Protocol for Improved Proteomic Analysis. Methods Mol Biol 2420:1–10.

57. Guerrero L, Carmona-Rodriguez L, Santos FM, Ciordia S, Stark L, Hierro L, Perez-Montero P, Vicent D, Corrales FJ. 2024. Molecular basis of progressive familial intrahepatic cholestasis 3. A proteomics study. Biofactors doi:10.1002/biof.2041.

58. Colome N, Abian J, Aloria K, Arizmendi JM, Barcelo-Batllori S, Braga-Lagache S, Burlet-Schiltz O, Carrascal M, Casal JI, Chicano-Galvez E, Chiva C, Clemente LF, Elortza F, Estanyol JM, Fernandez-Irigoyen J, Fernandez-Puente P, Fidalgo MJ, Froment C, Fuentes M, Fuentes-Almagro C, Gay M, Hainard A, Heller M, Hernandez ML, Ibarrola N, Iloro I, Kieselbach T, Lario A, Locard-Paulet M, Marina-Ramirez A, Martin L, Morato-Lopez E, Munoz J, Navajas R, Odena MA, Odriozola L, de Oliveira E, Paradela A, Pasquarello C, de Los Rios V, Ruiz-Romero C, Sabido E, Sanchez Del Pino M, Sancho J, Santamaria E, Schaeffer-Reiss C, Schneider J, de la Torre C, Valero ML, Vilaseca M, et al. 2022. Multi-laboratory experiment PME11 for the standardization of phosphoproteome analysis. J Proteomics 251:104409.

59. Zhou Y, Zhou B, Pache L, Chang M, Khodabakhshi AH, Tanaseichuk O, Benner C, Chanda SK. 2019. Metascape provides a biologist-oriented resource for the analysis of systems-level datasets. Nat Commun 10:1523.

60. Clarke DJB, Kuleshov MV, Schilder BM, Torre D, Duffy ME, Keenan AB, Lachmann A, Feldmann AS, Gundersen GW, Silverstein MC, Wang Z, Ma’ayan A. 2018. eXpression2Kinases (X2K) Web: linking expression signatures to upstream cell signaling networks. Nucleic Acids Res 46:W171–W179.

61. Chen EY, Xu H, Gordonov S, Lim MP, Perkins MH, Ma’ayan A. 2012. Expression2Kinases: mRNA profiling linked to multiple upstream regulatory layers. Bioinformatics 28:105–11.

62. Huang Q, Szklarczyk D, Wang M, Simonovic M, von Mering C. 2023. PaxDb 5.0: Curated Protein Quantification Data Suggests Adaptive Proteome Changes in Yeasts. Mol Cell Proteomics 22:100640.

63. Ciordia S, Santos FM, Dias JML, Lamas JR, Paradela A, Alvarez-Sola G, Avila MA, Corrales F. 2024. Refinement of paramagnetic bead-based digestion protocol for automatic sample preparation using an artificial neural network. Talanta 274:125988.

